# Re-convolving the compositional landscape of primary and recurrent glioblastoma reveals prognostic and targetable tissue states

**DOI:** 10.1101/2021.07.06.451295

**Authors:** Osama Al-Dalahmah, Michael G. Argenziano, Adithya Kannan, Aayushi Mahajan, Julia Furnari, Fahad Paryani, Deborah Boyett, Akshay Save, Nelson Humala, Fatima Khan, Juncheng Li, Hong Lu, Yu Sun, John F. Tuddenham, Alexander R. Goldberg, Athanassios Dovas, Matei A. Banu, Tejaswi Sudhakar, Erin Bush, Andrew B. Lassman, Guy M. McKhann, Brian J. A. Gill, Brett Youngerman, Michael B. Sisti, Jeffrey N. Bruce, Peter A. Sims, Vilas Menon, Peter Canoll

## Abstract

Glioblastoma (GBM) is an aggressive diffusely infiltrating neoplasm that spreads beyond surgical resection margins, where it intermingles with non-neoplastic brain cells. This complex microenvironment harboring infiltrating glioma and non-neoplastic brain cells is the origin of tumor recurrence. Thus, understanding the cellular and molecular features of the glioma microenvironment is therapeutically and prognostically important. We used single-nucleus RNA sequencing (snRNAseq) to determine the cellular composition and transcriptional states in primary and recurrent glioma and identified three compositional ‘tissue-states’ defined by the observed patterns of cohabitation between neoplastic and non-neoplastic brain cells. These comprise states enriched in A) neurons and non-neoplastic glia, B) reactive astrocytes and inflammatory cells, and C) proliferating tumor cells. The tissue states also showed distinct associations with the different transcriptional states of GBM cells. Spatial transcriptomics revealed that the cell-types/transcriptional-states associated with each tissue state colocalize in space. Tissue states are clinically significant because they correlate with radiographic, histopathologic, and prognostic features. Importantly, we found that our compositionally-defined tissue states are enriched in distinct metabolic pathways. One such pathway is fatty acid biosynthesis, which was enriched in tissue state B – a state enriched in recurrent glioblastoma and associated with shorter overall survival- and composed of astrocyte-like/mesenchymal glioma cells, reactive astrocytes, and monocyte-like myeloid cells. We showed that treating acute slices of GBM with a fatty acid synthesis inhibitor is sufficient to deplete the transcriptional signature of tissue state B. Our findings define a novel compositional approach to analyze glioma-infiltrated tissue which allows us to discover prognostic and targetable features, paving the way to new mechanistic and therapeutic discoveries.

## Introduction

Glioblastoma (GBM) is the most malignant glial tumor of the brain and is refractory to current treatment. Although gross surgical resection of the visible tumor is sometimes feasible, glioma cells infiltrate the brain beyond the resection margins. While many studies have characterized the transcriptional and genomic features of GBM cells and glioma associated microglia/myeloid cells, a comprehensive analysis of other cells in the GBM microenvironment, and the patterns of cohabitation of different cell types is lacking. Previous studies have shown that the composition of glioma infiltrated samples varies from cellular tumor comprised of GBM and myeloid cells, to minimally infiltrated GBM margin tissue composed largely of non-neoplastic brain microenvironment cells, including neurons and glia [1–3]. This is the microenvironment into which tumor cells migrate and proliferate, leading to recurrence, and is also the target of adjuvant therapy. Thus, understanding the cellular milieu of the tumor microenvironment at presentation and recurrence, including both neoplastic and non-neoplastic cells, is vital for advancing the management of GBM. Our goal is to determine patterns of cellular composition and transcriptional states in primary and recurrent GBM, including both neoplastic glioma cells and non-neoplastic brain cells.

Early studies used bulk RNA-sequencing approaches to understand GBM states in MRI-localized samples from contrast-enhancing (CE) and non-contrast enhancing (NCE) margins [3–6]. More resolution is attained using single cell RNAseq (scRNAseq) approaches, which are being increasingly used to understand heterogeneity in gliomas. Several studies have employed scRNAseq from freshly resected surgical samples to explore the heterogeneity of GBM [2, 7–12]. These studies have significantly advanced our understanding of the heterogeneity and pathology of glioma. However, application of whole cell scRNAseq is faced with practical challenges related to the limitations of acquiring and processing freshly resected glioma tissue and the technical incompatibility with banked frozen glioma tissue. Moreover, scRNAseq is limited in sampling non-neoplastic cells of the microenvironment like neurons and astrocytes, which are major constituents of the tumor-margins [2, 8–11], in part because of cell-type survivability/selection bias during tissue dissociation. Thus, while advances have been made in defining the genetic alterations in glioma [13, 14] and the transcriptional states of glioma cells and immune cells [15–18], comprehensive analyses of cellular composition and diversity of cellular phenotypes in primary and recurrent gliomas remain a challenge.

Here, we circumvented these limitations of scRNAseq by using single-nucleus RNA-sequencing (snRNAseq), which allowed us to analyze frozen tissue, and inclusively sample cells of the microenvironment from primary and recurrent glioma. We sampled glioma-infiltrated tissue from cellular tumor to minimally infiltrated surrounding brain tissue at the single cell level. Transcriptional analysis of copy number variations (CNVs) provided a metric to distinguish neoplastic (CNVpos) and non-neoplastic (CNVneg) nuclei, and unbiased clustering revealed that primary and recurrent tumors harbor CNVpos glioma cells with similar transcriptional states. Conversely, the microenvironment of primary and recurrent glioma displayed distinct cell-type specific states and different compositional landscapes. Leveraging information from the snRNAseq-derived compositional make-up of glioma-infiltrated samples defines three generalizable “tissue-states” with each tissue-state showing enrichment for specific gene signatures that can be identified in bulk RNAseq samples. These compositional patterns were also examined using spatial transcriptomics, which revealed colocalization of specific neoplastic and non-neoplastic cell types. We demonstrate that tissue-states are prognostically relevant and display metabolic dependencies that can be pharmacologically targeted.

## RESULTS

### Transcriptional analysis of the glioma microenvironment reveals prognostically significant subpopulations of non-neoplastic astrocytes

Given the importance of glioma microenvironment in tumor progression, we decided to investigate the implications of microenvironmental states on the prognosis of GBM. To achieve this, we first identified neoplastic and non-neoplastic nuclei based on chromosomal copy number variation (CNV) inference (Supplementary results). Based on the repertoire of transcriptional states of glioma cells that have been previously described ([2, 7–10, 12]), we confirmed that our CNV positive (CNVpos) neoplastic nuclei from primary and post-treatment recurrence GBM recapitulate known transcriptional states. We provide this data in the supplementary results including discussion of glioma states in primary and recurrent glioma (**Figures S1, S3**), CNV analysis of primary and recurrent glioma samples (**Figures S2, S4**), localization studies of glioma states in the tissue (**Figure S5**), and details on other low-grade glioma and epilepsy samples included in this study (**Figures S6-7**). We focused on the non-neoplastic CNV negative (CNVneg) nuclei of the glioma microenvironment and combined in our analysis nuclei from primary and recurrent glioma, as well as nuclei from low-grade glioma (LGG) and epilepsy, to include a spectrum of neurological diseases with alterations to non-neoplastic cells in the brain microenvironment. The clinical data on the samples, QC metrics, and number of nuclei per lineage/cluster is provided in **Supplementary Table-1.** Our CNVneg nuclei datasets included 16831 nuclei: 6929 from primary glioma, 6008 from post-treatment recurrent glioma, 2875 from epilepsy, and 1019 from LGG. We projected these nuclei in UMAP space and assigned cell lineages as shown in **Figure-1A**. The expression of a select number of marker genes per lineage is shown in **Figure-1B**. We present the results on myeloid lineage nuclei in the supplementary results (**Figure-S9**), which demonstrates that monocyte-derived tumor-associated macrophages (TAMs) were enriched in recurrent glioma, while microglia-derived TAMs were enriched in primary glioma, consistent with a previous report[15].

We focused on astrocytes, which are key elements of the glioma microenvironment and are not well represented in glioma single-cell RNAseq datasets [2, 7–12, 19]. A recent paper implicated GBM-associated astrocytes in promoting an immunosuppressive microenvironment [20]. Moreover, the distinction between tumor-astrocytes and reactive-astrocytes is of major diagnostic importance in neuropathology. Thus, we analyzed astrocytes (707 nuclei – 284 from primary glioma, 254 from recurrent glioma, 45 from LGG, and 121 from epilepsy) in isolation from other cell types, performed linear dimensionality reduction, and clustered them into three states; Ast1 – protoplasmic astrocytes, Ast2 – reactive astrocytes with expression of oligodendroglial and neuronal genes, and Ast3 – reactive astrocytes with inflammatory gene expression (**Figure-1C Supplementary Results, and Supplementary Table 4**). The astrocytes are projected by disease condition in **Figure 1D**. Clustering of astrocytes was based on the enrichment of three genesets with pre-defined genes relevant to astrocyte function (**Supplementary Table 4** and **Figure 1F)**. Expression of select markers of these astrocytes states (clusters) is shown in **Figure 1F**. Since astrocytes and glioma shared gene signatures (for example, CLU and LGALS3 expression), we performed differential gene expression analysis between primary and recurrent glioma non-neoplastic astrocytes and all CNVpos glioma nuclei and identified 1620 genes were higher in astrocytes compared to glioma and 3380 were higher glioma compared to astrocytes. Examples of genes higher in non-neoplastic astrocyte include genes associated with Alzheimer’s disease (CLU, APOE)[21, 22], metallothionein genes (MT1H, MT1G, MT1M, MT1F, MT1E, MT1X, MT2A, and MT3 – increased in reactive astrocytes [23]), Synuclein genes (SNCA, SNCB, and SNCG), WIF1, CHI3L2 (associated with poor prognosis in glioma[24]), ALDOC, ALDOA, AQP4, carbonic anhydrases CA2 and CA11, and CXCL14, a cytokine implicated in promoting glioma invasion [25] (**Supplementary Table-4)**. Conversely, genes higher in CNVpos glioma include EGFR, PTPRZ1, NOVA1, CD24, Nestin (NES), SOX5, and SOX4. We used KEGG pathway enrichment analysis to query the function of these genes (**Figure-1G-H**). Further analysis of the differentially expressed genes showed that several KEGG pathways were enriched in genes higher in non-neoplastic astrocytes (**Figure-1G)**, with some relating to neurodegeneration such as Parkinson disease, and prion disease. Notably, these signatures are highly enriched in oxidative phosphorylation genes (**Supplementary Table-4**), which are dysregulated in neurodegenerative diseases [26]. Moreover, other metabolic pathways enriched in astrocyte DEGs included metabolism of fatty acids, glycolysis, TCA cycle, and ferroptosis. Conversely, KEGG pathways increased in CNVpos tumor-astrocytes were largely related to DNA replication, cancer-related pathways including ErbB and MAPK signaling, DNA replication and mismatch repair (**Figure-1H** and **Supplementary Table-4)**.

**Figure 1:**
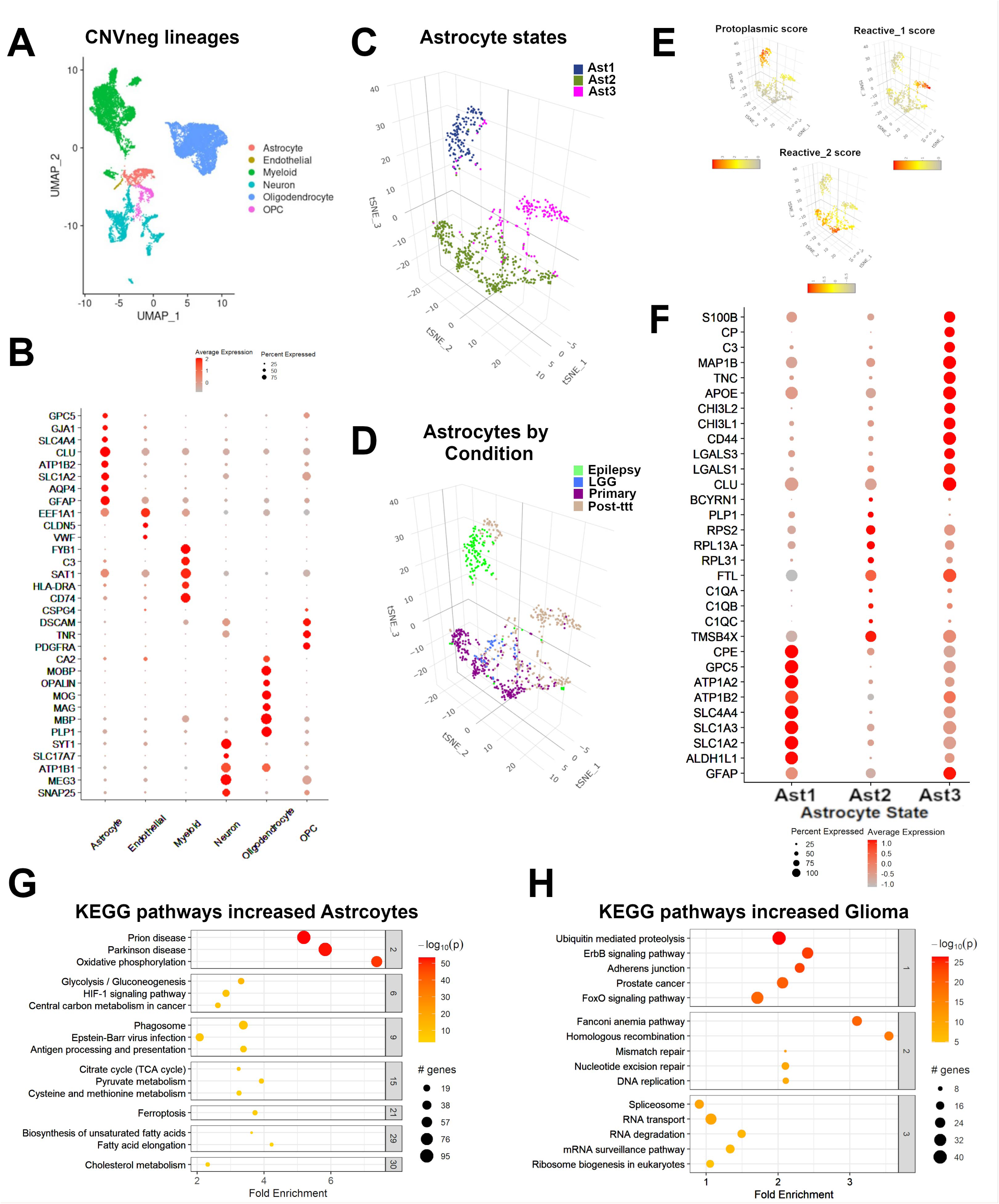
snRNAseq identifies three transcriptionally distinct astrocytes states in the glioma microenvironment. **A**) Uniform-manifold approximation and projection (UMAP) graphs showing putative non-neoplastic (CNVneg) from primary glioma, recurrent glioma, low grade glioma (LGG) - and epilepsy (see supplementary data for the analysis of LGG and epilepsy cases). The nuclei are color-coded by lineage (Oligodendrocytes, oligodendrocyte-precursor cells (OPC), neurons, astrocytes, myeloid cells, and endothelial cells). **B**) Dot plots showing normalized expression of select lineage genes (row) in the lineages from A (columns). The size of each circle corresponds to the proportion of each lineage that expresses a given gene. **C**) Three-dimensional tSNE plots showing all astrocyte nuclei color-coded by astrocyte state (Ast1 – protoplasmic astrocytes, Ast2 – reactive astrocytes with misexpression of non-astrocyte lineage genes, and Ast3 – reactive astrocytes with expression of inflammatory genes. **D**) Three-dimensional tSNE plots showing all astrocyte nuclei color-coded by disease condition. **E**) tSNE plots showing the enrichment of gene signatures used for astrocyte clustering in astrocyte nuclei. **F**) Gene expression dot plots showing select gene marker expression for the astrocyte states. **G-H**) Active subnetwork enrichment analysis of KEGG pathways in genes differentially expressed in CNVneg glioma-associated astrocytes compared to CNVpos glioma cells in primary and recurrent IDH-WT glioma. Fold enrichment is represented on the x-axis and the pathways in the y-axis. The pathways are clustered to denote shared genes driving enrichment. The size of the circle per pathway denotes the number of enriched genes, and the negative log10 of the adjusted p.value is represented by color. Pathways enriched in genes significantly higher in astrocytes compared to glioma cells are shown in **G** and include neurodegenerative diseases and oxidative phosphorylation, metabolism including fatty acid metabolism. Pathways enriched in genes significantly higher in glioma cells compared to astrocytes are shown on the **H** and include DNA replication, splicing, and ErbB signaling.

snRNAseq and spatial transcriptomics reveals patterns of co-habitation between neoplastic and non-neoplastic cell types

Given the heterogeneity of cellular states of glioma and non-neoplastic cells in the glioma microenvironment, we hypothesized that the transcriptional landscape of GBM is determined by patterns of cohabitation of specific types and transcriptional states of neoplastic and non-neoplastic cells. To test this hypothesis, we first asked if specific glioma, or brain microenvironment lineages were differentially abundant or deplete across primary and recurrent glioma using a regression model [27] to test for differential abundance (**Figure-2A**). The results showed that for CNVpos cells, gl_Mes2 were significantly more abundant in recurrent glioma, while gl_PN1 were more abundant in primary glioma, (Benjamini-Hochberg adjusted p values (q-value) 3.99e-2 and 1.318e-5 respectively). For the CNVneg cells in the glioma microenvironment, OPCs were significantly more abundant in primary glioma (q-value 1.085e-03). These results show that patterns of cellular composition vary in primary and recurrent glioma, and likely contribute to determining the transcriptional landscape of glioma.

To uncover patterns of ‘tissue-states” with correlated cell states/lineages, we took advantage of the relatively unbiased sampling of cellular composition in the brain tumor microenvironment provided by snRNAseq. We approximated the cellular composition of each surgical sample by recombining the cells from all the distinct cell populations, as identified by snRNAseq, to create a compositional matrix containing the abundance of all cell types across all samples (**Supplementary Table-1**). The cellular composition matrix includes three astrocytic clusters (Ast1-3), five immune-cell states (Myel1, moTAM, mgTAM, prTAM, and T cells - see supplementary results and methods), neurons, oligodendrocytes, endothelial cells, OPCs, and glioma cells. We then used principal component analysis of the resulting cellular composition matrix and identified the compositional features that account for the variance across the samples (**Figure-2B**). We used the glioma states as supplementary quantitative variables [28] – the coordinates of which can be predicted from the other variables input into the PCA analysis. The results showed that the relative abundance of CNVpos glioma cells versus CNVneg non-neoplastic cells (neurons, oligodendrocytes, OPCs) is the major feature of the first principal component, and the abundance of reactive astrocytes (Ast3), macrophage-like myeloid cells (moTAM), and T cells is the major feature of the second principal component. Notably, the abundance of a specific subpopulation of astrocyte-like/mesenchymal glioma cells (gl_Mes2) was also highly correlated with the second principal component (PC2). These findings indicate that specific subpopulations of neoplastic and non-neoplastic cells tend to co-inhabit glioma samples. To assess if the cohabitation of cell types and transcriptional states is prognostically relevant, we used the IDH-WT GBM TCGA and CGGA survival datasets and performed a log-rank test on samples with positive versus negative PC2 signature enrichment and found that positive enrichment is significantly associated with poor survival (**Figure-2C**). These data show that glioma infiltrated tissue shows patterns of cellular composition driven by co-habitation of specific cell-types and transcriptional states and reveal prognostically-relevant gene signatures that span across both neoplastic and non-neoplastic cell types.

To further characterize the cohabitation of specific cell types and transcriptional states in GBM, we analyzed nine samples of IDH-WT GBM infiltrated brain tissue using spatial transcriptomics (ST - **Supplementary Table-1**, **Figure 2D-E** and **Figures S10 and S11**). We deconvolved the ST data using RCTD[29] and analyzed the spatial relationships between cell types. To improve the accuracy of deconvolution results, we incorporated snRNAseq from the same tissue samples used to generate the ST data when possible (Validation snRNAseq dataset - see supplementary results and **Figure S8**. We determined the proportion of our 18 cell types present in each of the 9,017 transcriptomic capture spots that comprised our ST experiments and quantified the relationship between different cell types using spatial cross-correlation. Spatial cross-correlation quantifies the correlation between the proportion of cell type A in any given spot and the proportion of cell type B in that spot’s neighbors. By evaluating this metric for every spot, between all pairwise comparisons of cell types, we were able to quantitatively assess the geographic relationship between cell types and determine which global patterns of cohabitation were statistically significant (**Figure 2E)**. Clustering of the spatial cross-correlations showed three main clusters: 1-showing positive and statistically significant cross-correlations between non-neoplastic cell types such as neurons, oligodendrocytes, non-reactive astrocytes (Ast1, Ast2), and OPCs; 2-showing statistically significant spatial correlations between gl_Mes2, reactive astrocytes (Ast3), moTAM, and T-cells; and 3-showing statistically significant spatial correlations between endothelial cells and several CNVpos glioma cell subtypes. Interestingly, gl_PN2 and gl_Mes2 were significantly spatially cross-correlated, and our independent validation dataset analyzed by RNAscope for gl_Mes2 and gl_PN2 markers (**Figure S5**) shows both are significantly more abundant in cortical regions. These findings provide additional evidence to support cohabitation of these cell types.

**Figure 2:**
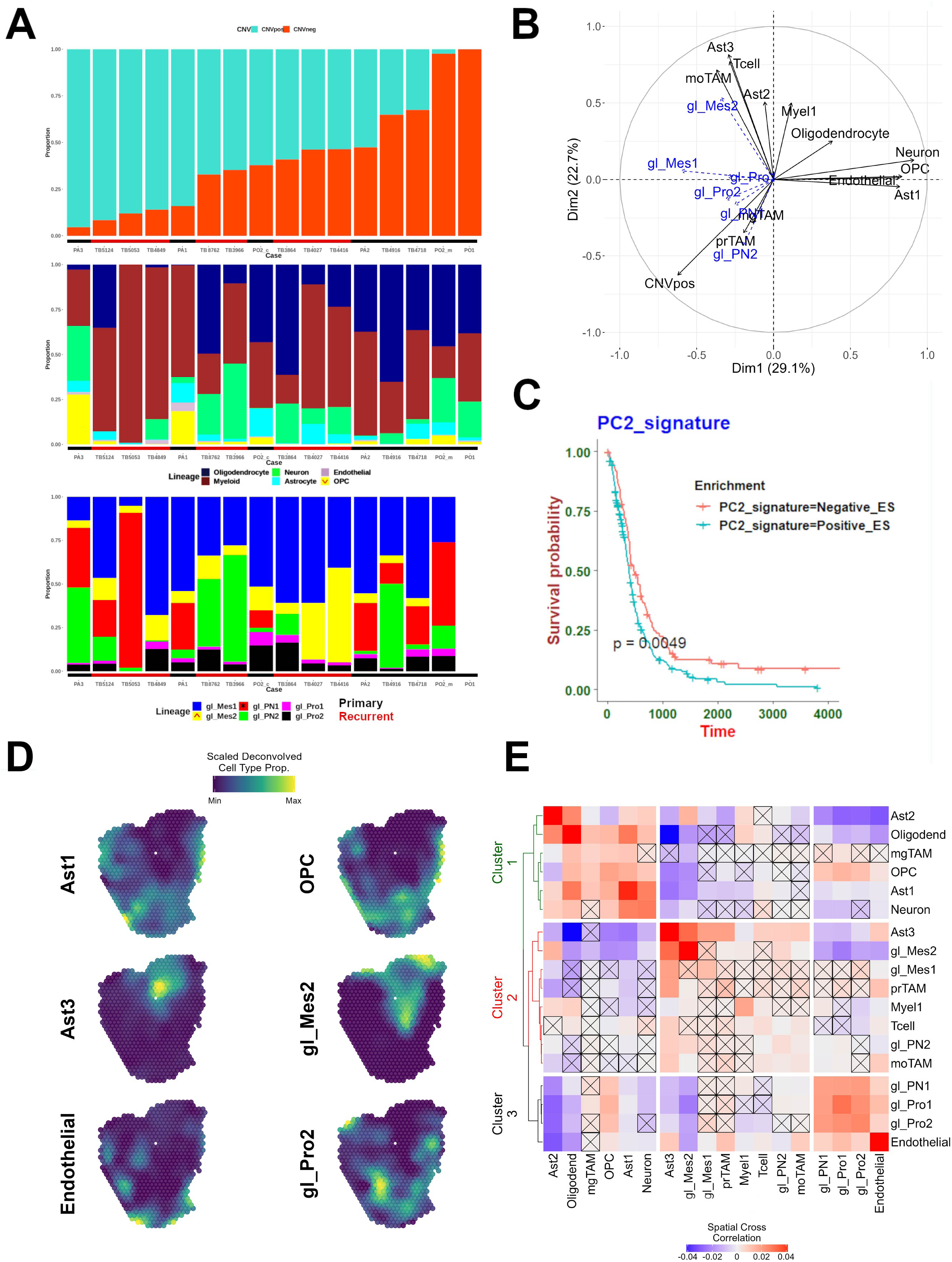
snRNAseq and spatial transcriptomics identify patterns of co-habitation that correlate with survival. **A**) Bar plots demonstrating the fractional composition of each one of 16 samples analyzed by snRNAseq (8 primary IDH-WT glioma from 7 patients, one case was divided to core and overlying cortex, and 8 recurrent IDH-WT glioblastoma) The first row of bar plots represent the fraction of neoplastic (CNVpos) and non-neoplastic (CNVneg) nuclei. The middle row represents the fraction of the non-neoplastic nuclei contributed by neurons, oligodendrocytes, OPCs, astrocytes, myeloid cells, endothelial cells, and astrocytes. The red arrowhead in the OPC legend box indicate that OPCs were significantly reduced in recurrent GBM as determined by differential abundance analysis. The bottom row represents the fraction of the neoplastic nuclei contributed by proneural/progenitor-like glioma (gl_PN1, gl_PN2), astrocyte-like/mesenchymal glioma (gl_Mes1, gl_Mes2), and proliferative glioma (gl_Pro1, and gl_Pro2). The description of glioma states is provided in the supplementary results. The red arrowhead in the gl_Mes2 legend box and the black star in gl_PN1 indicate that gl_Mes2 and gl_PN1 were significantly increased and reduced in recurrent GBM, respectively, as determined by differential abundance analysis. **B**) Principal component analysis of the fractional composition matrix of 19 samples encompassing eight primary and eight recurrent gliomas plus three epilepsy samples. The tissue composition matrix consists of the percentage of nuclei per each tissue state. Immune cell states are: mgTAMs (microglia-derived Tumor-associated macrophages), moTAM (monocyte-derived TAMs), prTAM (proliferative TAM), Myel1 (baseline myeloid cells), and T cells. Astrocyte states include baseline (protoplasmic) astrocytes (Ast1), reactive CD44+ astrocytes (Ast3), and reactive astrocytes with expression of non-astrocyte genes (Ast2) – see text and supplementary results for additional description of these cell states. CNVpos represents the total percentage of all tumor states per sample. Individual tumor states were not used in PCA calculation, rather they were used as used supplementary quantitative variables and their coordinates were predicted from the PCA analysis – see methods. **C**) Kaplan-Meier survival plot graphing survival in the combined TCGA and CGGA RNAseq datasets. The samples were classified based on enrichment of gene signatures of microenvironment states correlated with PC2 into positive or negative enrichment. Statistical significance was computed using the log rank test. **D**) Representative plots showing deconvolved proportions of select cell types and glioma states across an ST sample. Each subplot is range scaled for the proportion of that cell type in the sample to show the relative spatial distribution of that cell type. **E**) Heatmap showing the average spatial cross-correlation between all cell types at a radius of 900µm surrounding spatial transcriptomic spots across all nine ST experiments. The diagonal of the matrix, which shows the spatial cross-correlation between a cell type and itself, is an indication of the degree of spatial autocorrelation in that cell type and is not necessarily equal to one. Spatial cross-correlation relationships were tested for significance using permutation (see methods), and non-significant relationships are denoted with an X. Hierarchical clustering of the distance matrix derived from the cross-correlation matrix produced three clusters.

### Re-convolution of snRNAseq identifies three tissue states based on cellular composition of glioma and its microenvironment

Driven by the above findings, we clustered the snRNAseq samples from our discovery dataset into 3 distinct “tissue-states” based on the approximated cellular compositions described above; **tissue-state A** samples are predominantly composed of non-neoplastic brain cells, including neurons oligodendrocytes, and OPCs**, tissue-state B** samples are enriched in reactive astrocytes, myeloid/macrophages, and T-cells, and **tissue-state C** samples are predominantly composed of CNVpos glioma cells (**Figure 3A-B**). Based on the results of our compositional clusters/tissue states, we are able to assign tissue states to an external validation set based on k-means classification (validation set – **supplement S8H**). To generate a gene signature for each tissue state, we combined the snRNAseq for all nuclei in each sample and performed differential gene expression analysis between tissue-state clusters, using the pseudobulk expression profile of each sample as a biological replicate. This analysis identified the top-differentially expressed genes unique to each tissue state (**Supplementary Table-7**). To assess the generalizability of the three tissue-states, we performed single sample GSEA analysis for the tissue state gene signatures using a dataset of bulk RNAseq analysis performed on 91 primary and recurrent MRI-localized samples from 39 patients. We found that these samples separated into 3 compositional clusters based on their enrichment score for snRNAseq-defined “tissue-states” (**Figure 3C**). We refer to the compositional clusters and tissue-states interchangeably henceforth. These three tissue-states are further demonstrated by projecting the RNA-expression levels for canonical markers of the predominant cell types for each tissue-state in **Figure 3E** showing RBFOX3 (neuronal marker) in tissue-state A, CD68 (myeloid marker) in tissue-state B, and MKI67 (proliferation marker) in tissue-state C. SOX2 (a pan-glioma marker) was widely distributed across the samples, indicating variable degrees of tumor infiltration across samples in all three tissue-states (**Figure 3E**). Further analysis revealed that these tissue-state gene signatures are enriched for specific biologically relevant functional ontologies. For example, tissue-state A is enriched of genes involved in synaptic transmission, tissue-state B is enriched for genes associated with inflammation, and tissue state C is associated with cell proliferation (**Figure 3D**). To further validate these findings, we quantified total cellularity and the IHC labeling indices SOX2, NeuN, CD68, and Ki67 in 45 recurrent and primary glioma samples (**Figure 3F**) and found that total cellularity was highest in cluster C, which also had the highest abundance of SOX2+ and Ki67+ cells, while cluster A had the highest abundance of NeuN+ cells, and Cluster B had the highest abundance of CD68+ cells. While Clusters A and B resemble normal and reactive brain tissue, the SOX2 and Ki67 labeling indices indicate that these clusters comprise samples with variable levels of glioma infiltration.

**Figure 3:**
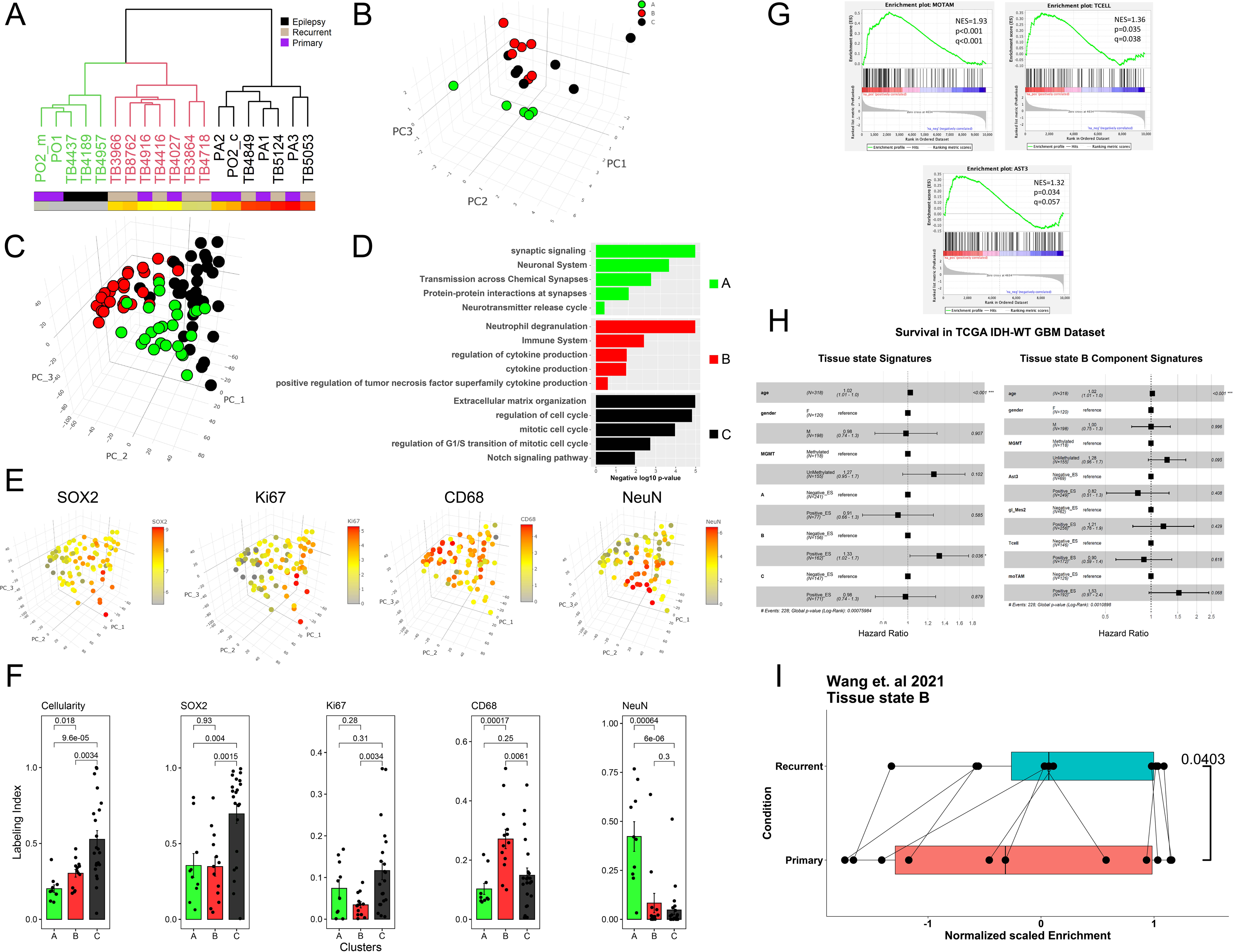
Tissue composition analysis defines “tissue states” recapitulated in validation a bulk RNAseq dataset that. **A**) Dendrogram of hierarchically clustered glioma and epilepsy samples based on Manhattan sample distance analysis drawn from the fractional composition matrix (see Figure 2A). Three clusters were identified and are color-coded on the dendrogram in black (Tissue-state C), red (Tissue-state B), and green (Tissue-state A). The condition (primary, recurrent and epilepsy) is indicated in the top bar underneath. The proportion of neoplastic nuclei (CNVprop) is indicated in the bottom bar. **B**) Three-dimensional scatter plot showing the samples in **A** projected in the first three principal component loadings – see figure 2B for PCA analysis. The samples are color-coded by cluster designation as in A. **C**) Bulk RNAseq samples from 91 primary and recurrent IDH-WT glioblastoma samples projected in the principal component space. The samples were clustered (Hierarchical clustering – Ward.D2 method) on the Euclidian distance of the enrichment scores of the genes unique to each tissue state signature into three clusters A-C and are color-coded as such. **D**) Gene ontology term analysis of the genes uniquely and differentially expressed in each of the clusters in **C**. The bar plots are color coded as per the clusters in C. KEGG, REACTOME, or Biological Process GO pathways are shown in the y-axis. Negative log10 of the adjusted p-value is shown on the x-axis. **E**) Normalized expression of select genes characteristic of each of the clusters projected onto the compositional-signature enrichment score space shown in C. Red denotes high expression, and grey denotes low expression. NeuN (RBFOX3) is highest in the samples of Cluster A. CD68 is highest in the samples of cluster B. SOX2 is highest in the samples of clusters B and C. MKI67 is highest in the samples of cluster C. **F**) Quantification of histological cellularity analysis (far left) as well as immunohistochemistry labeling indices of (from left to right) SOX2, KI67, CD68, and NeuN. The labeling index is shown on the y-axis. Note that the y-axis for the cellularity graph is total cellularity normalized to the most cellular sample. The sample clusters are labeled (A-C) as in **C**. p values were calculated using Kruskal-Wallis test and are indicated on the graphs. N=8 for cluster A, 25 for cluster B, and 12 for cluster C. **G**) Pre-ranked Gene Set Enrichment Analysis (GSEA) comparing tissue state B bulk RNAseq samples with tissue states A & C samples for 3 sub-lineages: Ast3, moTAM, and T-cells. Marker genes for each cell type were used as the gene set for each analysis. Normalized Enrichment Score (NES) is displayed, along with p-values and FDR-adjusted q-values. **H**) Cox proportional hazard ratio of survival in the combined TCGA and CGGA IDH-WT GBM dataset given enrichment of each of the tissue state signatures (left) and for the individual cell types that comprise Tissue State B (right). Age, sex, and MGMT status are included as co-variates in the model. The p values are shown on the left, bars indicate confidence intervals (also noted on the right). Enrichment of each geneset was categorized as negative or positive. **I**) Boxplots of the tissue state B normalized enrichment scores in the Wang, L. et al. 2021 paired primary and recurrent GBM dataset. Paired samples are denoted by connected points. Paired t-test – one-tailed. N= 11 per group. The p value is indicated.

To substantiate the clinical relevance of investigating glioma tissue in terms of tissue states, we investigated whether the enrichment of tissue state signatures correlated with survival in the TCGA-CGGA IDH-WT glioblastoma dataset. Given that tissue state B was enriched for the gene signatures of Ast3, moTAM, and T-cells (**Figure-3G**), and considering our findings in Figure-2C, we expected it to be associated with increased risk of death in survival cohorts. As expected, enrichment of Cluster B gene signature in the IDH-WT TCGA and CGGA datasets was associated with a significant increase in the hazard of death in cox proportional hazard regression model, with covariates controlled for including age, sex, and MGMT methylation status (**Figure 3H**). In contrast, no significant association with survival was seen for the gene signatures of the individual cell types that compose tissue state B, including gl_Mes2, Ast3, T-cells, and moTAMs (**Figure 3H).** To further establish the clinical relevance of taking a tissue state approach in investigating glioblastoma transcriptomics, we asked if the tissue state B differentially enriched in primary versus recurrent glioblastoma status. This question was especially relevant given that compositional cluster B was largely composed of recurrent glioma samples (**Figure-3A**). We thus asked if that signature is positively enriched in RNAseq profiles from previously published paired primary and recurrent glioblastoma samples [30] (**Figure-3I**). The results showed significant enrichment in tissue state B signatures in the recurrent GBM samples. Together, the results show that tissue state B signature is prognostic and enriched during GBM recurrence.

### Glioma-associated tissue states are targetable and associated with distinct metabolic states

Given the distinct cohabitation patterns that drive tissue states, we hypothesized that these patterns of cellular cohabitation are associated with metabolic dependencies. To test this hypothesis, we investigated whether metabolic pathways are differentially enriched in genes differentially expressed between bulk RNAseq samples of the three tissue states. Unbiased analysis of enrichment of KEGG pathways in genes differentially expressed between compositional clusters/tissue-states revealed that they exhibit enrichment of multiple unique and specific pathways (**Figure 4A**). Interestingly, several of the tissue state-enriched pathways were metabolic pathways. Tissue-state A showed highest enrichment for oxidative phosphorylation and beta-glutamate metabolism, tissue-state C was most enriched for pyrimidine, folate, and purine metabolism, and tissue-state B showed highest enrichment of fatty acid and lipid metabolism (**Figure 4B**). We focused on fatty acid biosynthesis genes, a tissue state B enriched pathway, and projected the average normalized expression per lineage as a heatmap in **Figure 4C**. We found that genes in this pathway were distributed across multiple cell types, suggesting that the metabolic status of a tissue can have distinct, but functionally related effects on different cell types in that tissue. Notably, fatty acid synthase (FASN), a rate-limiting enzyme in fatty acid synthesis[31], was most highly expressed in astrocytes and glioma cells **(Figure 4C).** FASN inhibition has been shown to kill glioma cells [32], however, the impact of FASN blockade on the glioma microenvironment is yet to be fully explored. Defining the effects of FASN blockade on the glioma microenvironment is important because fatty acid metabolism is a physiologic pathway that involves interactions between multiple cell types that reside in the same habitat. In non-neoplastic brain tissue, fatty acids are synthesized by astrocytes and are distributed to other cells including neurons and oligodendrocytes [31], where they drive physiologic and cellular functions like neuronal maturation, membrane synthesis [33], and neuroprotection [34]. We thus hypothesized blocking fatty acid synthesis pathway would interfere with the cells that make up tissue state B and/or their interactions, and therefore would lead to depletion of tissue state B signature in glioblastoma infiltrated brain. To test this hypothesis, we treated astrocytes and explants of human IDH-WT glioblastoma with the FASN inhibitor Cerulenin (5mg/ml) and measured gene expression using the plate-seq RNAseq (**Figure 4D**). Astrocytes treated with Cerulenin exhibited numerous differentially expressed genes compared with DMSO controls (**Supplementary table-6, Figure 4E**). Genes increased in treated astrocytes were enriched in KEGG and Reactome pathways involved in mTOR signaling, ferroptosis, and unfolded protein response, while those decreased in treated astrocytes were enriched in pathways involved in cell cycle. Cerulenin treatment did not alter astrocyte viability (data not shown). We then treated IDH-WT glioblastoma explants with DMSO or Cerulenin **(Supplementary table-6)** and measured gene expression (**Figure 4F**). We found that genes increased in Cerulenin treated astrocytes were significantly enriched in Cerulenin treated IDH-WT glioblastoma explants, and that the tissue state B signature was depleted (negatively enriched). It is important to note that the tissue state B gene signature used in this enrichment analysis does not contain any genes that are part of the fatty acid synthesis pathway gene ontology (**supplementary table-7**). Overall, these results demonstrate that tissue-states exhibit enrichment of metabolic pathways, which can be targeted leveraging compositional information and metabolic dependencies.

**Figure 4:**
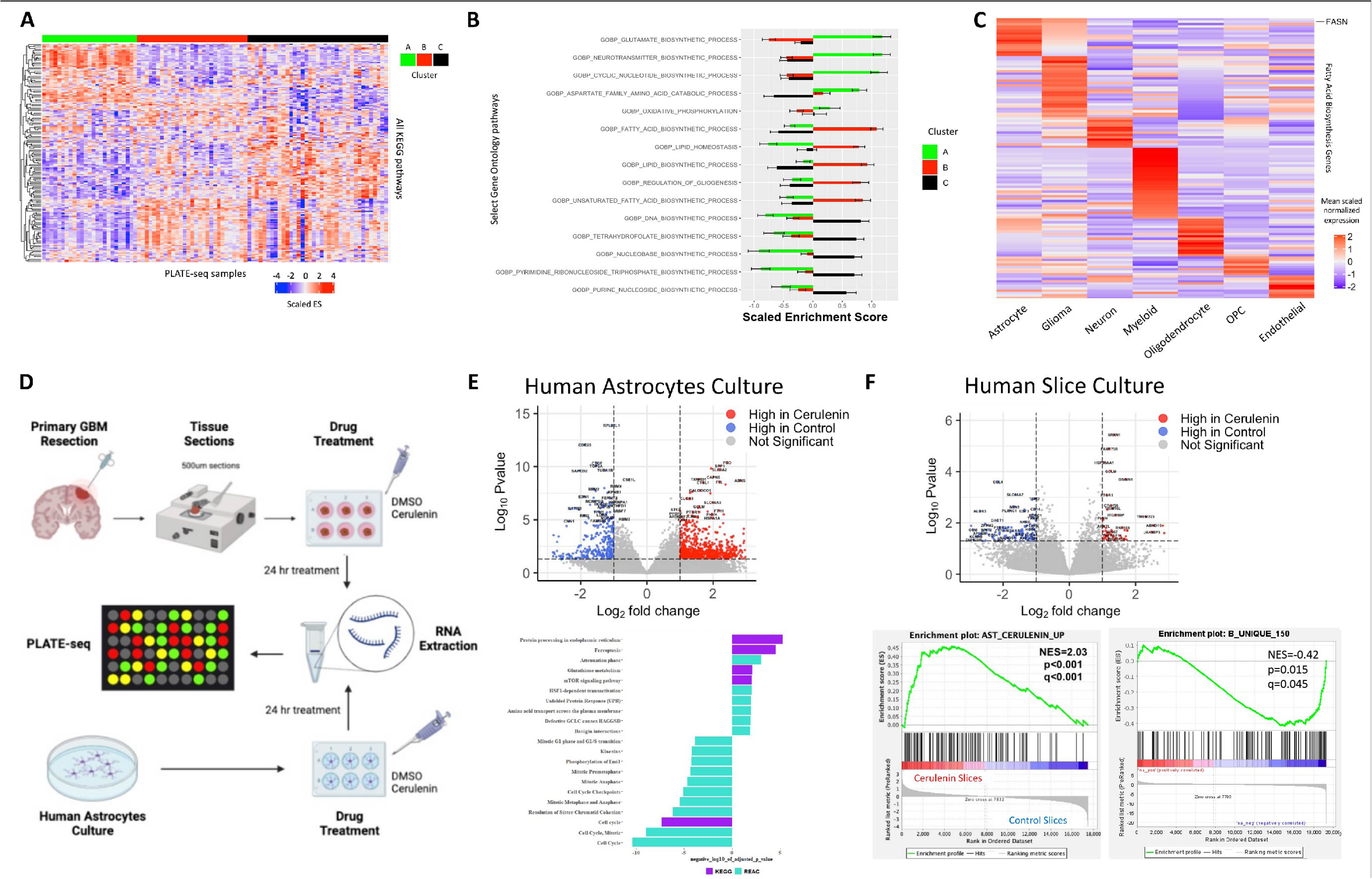
Metabolic pathways drive targetable tissue state signatures. **A**) Heatmap displaying scaled enrichment scores for all KEGG pathways across all PLATE-seq samples. The heatmap is grouped by tissue state (cluster A, B, C), annotated by the horizontal bar at the top. Hierarchical clustering was performed on the rows (pathways), demonstrating cluster-specific metabolic programs. **B**) Bar plot displaying scaled ssGSEA scores for select KEGG metabolic programs from A. Bar plots represent mean scaled ssGSEA score ± standard error for each of the three clusters for a given pathway. **C**) Representative example showing a heatmap displaying mean lineage-specific scaled normalized expression of genes in the GO: Biological Process - Fatty Acid Biosynthesis gene set – which was most enriched in tissue state B. Note the expression of the rate-limiting enzyme FASN is highest in astrocytes and glioma cells. **D**) Scheme of *in vitro* and *ex vivo* FASN perturbation studies. **E**) Volcano plot showing the log2 fold change (x-axis) and log10 p value (y-axis) of differentially expressed genes in astrocytes treated with Cerulenin (5mg/ml) versus control – Upper panel. Lower panel shows KEGG and Reactome pathway enrichment analysis with the terms indicated on the y-axis, and the log10 p value on the x-axis. The sign of the log10 p value indicates the direction of change (i.e. negative = reduced in Cerulenin treatment). **F**) Volcano plot showing the log2 fold change (x-axis) and log10 p value (y-axis) of differentially expressed genes in GBM slice cultures treated with Cerulenin (5mg/ml) versus control – Upper panel. Lower panel indicate GSEA plots of pre-ranked enrichment of the genes increased in astrocytes treated with Cerulenin (left) and the top 150 genes unique to tissue state B signature (right). The normalized enrichment scores (NES), p value (p), and adjusted p value (q) are indicated.

## DISCUSSION

In this work, we investigated the landscape of cellular composition and transcriptional states of neoplastic and non-neoplastic cell types in primary and post-treatment recurrent IDH-WT GBM using snRNAseq and spatial transcriptomics. Understanding heterogeneity in GBM is important for guiding treatment and meeting the challenge of recurrence. Recent studies revealed a diversity of glioma states that resemble cell lineages found during development and adulthood [2, 9–12, 19]. Our study provides a comprehensive analysis of the GBM microenvironment, including non-neoplastic cell types that are sparsely represented in datasets from prior studies using scRNAseq. Using a compositional approach rooted in relatively unbiased sampling of different GBM microenvironment cell types, we discovered that specific cell types/transcriptional states colocalize in “tissue-states”. Leveraging insight into correlated cellular states and lineages that co-inhabit tissue samples, we identified gene signatures that classify primary and recurrent GBM tissue into three tissue states: (A) normal brain, (B) reactive/inflammatory tissue, and (C) cellular/proliferative tumor. The tissue states exhibited variable levels of infiltrated by glioma cells. We stress that the tissue states we identified do not encompass the entirety of heterogeneity of possible tissue states, and discovery of new tissue states, for example in the context of different treatment scenarios, is highly probable. That said, the patterns of co-habitation in the tissue state model are further supported by deconvolution of spatial transcriptomics data, which highlights the differential distribution of specific neoplastic and non-neoplastic cell types.

Spatial cross-correlation analysis of our spatial transcriptomics data shows that transcriptionally distinct cell types exhibit significant patterns of colocalization. The patterns we observed using a neighborhood of 900 uM were similar to the tissue state patterns we identified using principal component analysis, demonstrating that these cell composition patterns can be observed using multiple approaches. Analysis of different neighborhood sizes may yield further insight into the mechanism that drive cohabitation between cell types. Patterns of cohabitation that span large areas are more likely to reflect environmental influences that can impact multiple cell types in a large geographic swathe, such as hypoxia or other metabolic stresses, whereas patterns that vary over smaller distances may reflect the effects of direct cell-cell interactions. Further investigation will be needed to elucidate the exact mechanisms that underlie the organization of cell types into distinct compositional patterns.

Importantly, we discovered that enrichment for tissue state B, a reactive state that harbors a reactive astrocyte state (Ast3), was associated with increased risk of death. The presence of tissue-state B was significantly associated with a worse mortality even though its individual component cell types were not, suggesting that the compositional patterns defined by tissue-states contribute to mortality in GBM. We show that gene signatures for these tissue states can also be identified in more accessible bulk RNAseq samples and correlate with immunohistochemical profiles. Significantly, we found that tissues states were transcriptionally enriched in distinct metabolic pathways, and that targeting fatty acid synthesis, a pathway enriched tissue state B, resulted in depletion of that signature in *ex vivo* GBM slice cultures. The therapeutic implications of our findings help expand the target of therapy from targeting one gene or one cell type, to targeting tissue states comprising cell populations that co-inhabit the tissue under defined metabolic constraints.

Our analysis of the cellular phenotypes in the glioma microenvironment revealed that subpopulations of non-neoplastic astrocytes show enrichment for abnormal transcriptional signatures that are also seen in the context of neurodegenerative diseases. In contrast to CNVpos neoplastic astrocytes, which express high levels of proliferation and glioma genes, a subpopulation of non-neoplastic astrocytes (Ast3) displayed a reactive signature reminiscent of astrocytes described in neurodegenerative diseases like Huntington disease, Parkinson disease and Alzheimer’s disease [23, 35]. This phenotype includes enrichment of pathways related to fatty acid metabolism. CLU, an astrocyte-expressed apolipoprotein involved in lipid transport [36] and neuroprotection in Alzheimer’s disease [21, 37], was significantly increased in glioma-associated astrocytes and is a marker of the reactive Ast3. We found that Ast3-like CLU-overexpressing astrocytes alter the transcriptional phenotype of glioma *in vitro* (**supplementary results and Figure S13**), including upregulation of genes involved in glial differentiation and notch signaling. Thus, our results point to commonalities in astrocyte dysregulation across neurologic diseases, which may offer therapeutic targets to be exploited in different clinical scenarios. Future studies are needed to further evaluate the potential of targeting reactive astrocytes as a therapeutic strategy to block GBM progression.

One of the main findings highlighted by our analysis of cellular composition is that specific cell types are correlated with each other both compositionally and spatially, indicating that they co-inhabit the same tissue-states. Cohabitation between cell types and transcriptional states was reflected in enrichment of distinct metabolic pathways. For example, tissue state B was enriched in the glutathione pathway, which determines a cell’s sensitivity to ferroptosis-inducing drugs [38], and in fatty acid metabolism, which has been implicated in glioma survival, stemness and progression [32, 39, 40]. We found that fatty acid metabolism genes were distributed among different cell types in the brain, however, FASN, the rate-limiting enzyme in fatty acid synthesis [31] was most highly expressed in astrocytes and glioma cells. Astrocytes play key roles in lipid metabolism; for example, in synthesizing fatty acids necessary for neuronal membranes [33] and catabolizing fatty acids released by neurons during excitotoxicity [41]. We showed that blocking FASN effectively depleted tissue state B signature from treated GBM slices. This may be explained by either a change in the composition of the GBM slices, given that FASN inhibition may lead to glioma cell death [32, 40], a change of gene expression of the cells that reside in the slices, or both. The latter is likely the case, given that GBM slices treated with FASN inhibitor showed a positive enrichment for the gene signature of astrocytes treated with FASN inhibitor, and negative enrichment for tissue state B. These finding are clinically relevant, given FASN is a promising target against glioblastoma [42], and highlight how the tissue-state approach can provide new insights into the effects of targeted therapies on the GBM microenvironment.

This study showed that the concept of tissue-states based on cellular cohabitation patterns generates testable hypotheses that inform our understanding of GBM biology. We found that tissue state B, which is enriched in reactive astrocytes (Ast3), monocyte-like tumor-associated myeloid cells, T-cells, and mesenchymal/astrocyte-like GBM cells (gl_Mes2) is associated with worse prognosis in GBM. Tissue state B is also characterized by specific metabolic signatures, like fatty acid metabolism, which we targeted *ex vivo* and showed it depleted the tissue state B signature. Future studies are needed to further evaluate the potential of targeting fatty acid synthesis to block GBM progression.

## Methods

### Human subjects and glioma tissue

Frozen primary untreated GBM tissue was acquired from the Bartoli brain tumor bank at Columbia University Medical Center. All diagnoses were rendered by specialized neuropathologists. Study protocols were approved by Columbia University Medical Center Institutional Review Board. All clinical samples were de-identified prior to analysis. Analyses were carried out in alignment with the principles outlined in the WMA Declaration of Helsinki and the Department of Health and Human services Belmont Report. Informed written consent was provided by all patients. The demographics of the cases used are provided in **Supplementary Table 1**.

### Extraction of nuclei and snRNAseq procedure

Nuclei were isolated from frozen surgical resection specimen as described in Al-Dalahmah O et al. 2020. Briefly, the frozen tissue samples were dissected from fresh frozen tissue or frozen OCT-embedded tissue blocks to yield tissue measuring in general from 5 x 2 x 1 mm to 10 x 6 x 3 mm. The tissue was homogenized using a dounce homogenizer in ice-cold 30% sucrose 0.1% Triton-X 100 based homogenization buffer. 10-15 strokes of the loose dounce pestle were followed by 10-15 strokes of the tight dounce pestle on ice. Mixing using a P1000 pipette followed before filtration through a BD Falcon 40um filters. Filtration was repeated after a 10-minute spin at 1000g at 4c. A cleanup step followed using a density gradient step as described in [43]. The nuclear pellet was suspended in 1% BSA in PBS resuspension buffer containing RNAse inhibitors. A final filtration step using 20um Flowmi ™ filters followed before dilution to 700-1200 nuclei per ul in resuspension buffer. The nuclear suspensions were processed by the Chromium Controller (10x Genomics) using single Cell 3’ Reagent Kit v2 or v3 (Chromium Single Cell 3’ Library & Gel Bead Kit v2, catalog number: 120237; Chromium Single Cell A Chip Kit, 48 runs, catalog number: 120236; 10x Genomics).

### Sequencing and raw data analysis

Sequencing of the resultant libraries was done on Illumina NOVAseq 6000 platformV4 150bp paired end reads. Alignment was done using the CellRanger pipeline (10X Genomics) to GRCh38.p12 (refdata-cellranger-GRCh38-1.2.0 file provided by 10x genomics). Count matrices were generated from BAM files using default parameters of the DropEst pipeline (Petukhov V et al. 2018). Filtering and QC was done using the scater package (3). Nuclei with percent exonic reads from all reads in the range of 25-75% were included. Nuclei with percent mitochondrial reads aligning to mitochondria genes of more than 19% were excluded. Genes were filtered by keeping features with >10 counts per row in at least in 31 cells. Further filtering of low quality cells was done to include cells with at least 400 detected genes and 10,000 reads.

### Single Nuclei RNAseq analysis

#### Sequencing and analysis of raw data

Sequencing of the resultant libraries was done on Illumina NOVAseq 6000 platformV4 150bp paired end reads. We used 10X chromium v2 chemistry for samples PO1 and PO2, and v3 chemistry for samples PA1, PA2, and P3. Read alignment was done using the CellRanger pipeline (v3.1 - 10X genomics) to reference GRCh38.p12 (refdata-cellranger-GRCh38-1.2.0 file provided by 10x genomics). Count matrices were generated from BAM files using default parameters of the DropEst pipeline [44].

#### Data-cleanup

Filtering and QC was done using the scater package [45, 46]. Nuclei with percent exonic reads from all reads in the range of 25-73% were included. Nuclei with percent mitochondrial reads aligning to mitochondria genes of more than 15% were excluded. Genes were filtered by keeping features with >10 counts per row in at least in 31 cells. The count matrix of each sample was normalized by first running the quickcluster function, then estimating size-factors by calling scran::computeSumFactors() function with default options and clusters set to clusters identified by calling quickcluster function. scater::normalize() function was then used to generated normalized counts. Doublet identification was done using scran::doubletCells function with default options, and cells with doublet score of NMADs > 3 were excluded as we described previously [23].

#### Combining multiple datasets from different sequencing batches

To control sequencing and technical batches, we utilized canonical correlation analysis in Seurat [47] accounting for batch and mitochondrial read percentage for CNVneg nuclei. For CNVpos nuclei, we accounted for case and mitochondrial read percentage.

#### Pre-Clustering and clustering of nuclei

Pre-clustering of nuclei was done in Seurat using the shared nearest neighbor smart local moving algorithm. PCA reduction was used as the reduction in the FindNeighbors() step. Pre-cluster identity determination was done using geneset enrichment analysis of lineage markers [23] and by inspecting cluster markers generated by scran::findmarkers(direction=”up”) function. Microglia +/- oligodendrocytes were used as negative control cell for InferCNV pipeline (below). Once CNVneg cells were verified, cells from all cases were aligned using Seurat and clustered. Clusters with mixed identities based on enrichment of multiple lineage genes were sub-clustered iteratively until all “pre-clusters” showed pure identities. Only then do we combine the pre-clusters of the same lineage into lineages (Astrocytes, neurons, oligodendrocytes, myeloid, endothelial). For subclustering of astrocytes and myeloid cells, we analyzed the nuclei in isolation of other lineages, and re-aligned them in Seurat, and reduced the dimensions before subclustering. For CNVpos nuclei, unbiased clusters were combined into glioma states/lineages based on similarity in marker expression and enrichment for known gene sets described in **Figures S1D and S3D**.

#### Count normalization

Raw counts were normalized in Seurat using the sctransform function SCT() function with default settings and controlling for percent mitochondrial gene expression [48].

#### Copy number variation analysis of snRNAseq

To detect putative neoplastic tumor cells, we used combination of marker expression and large scale copy number variation inference as per the InferCNV R package [49]. We used the default parameter as described in the package documentation. As a control population, we used microglia and oligodendrocytes from case PO2_1. Iteratively, CNVneg clusters including Oligodendrocytes and Neurons were identified and added as control cells. Different gene window sizes were tested (50, 100, 200) and yield similar results. We then applied an orthogonal approach to label putative neoplastic cells based on previous approaches described in [2, 50]. Briefly, Log2+1 counts were averaged across chromosomes for each nucleus. A principal component analysis (PCA) was performed on autosomal chromosomes in factominer R package [28]. Chromosomes with high correlation with PC2 were the same as the ones shown in the detected by inferCNV() with the exception is PO1, where no neoplastic cells were detected by InferCNV or CONICS. A malignancy score was calculated by dividing the log2 gained chromosome counts over the sum of those that are lost (selection was limited to three chromosomes or less). The scores were then z-neoplastic per sample. To identify putative neoplastic nuclei in this method, we next performed K means clustering of the scaled malignancy scores in R using the kmeans function and centers argument set to 2. PO1 does not show bimodal malignancy score distribution and the results of kmeans clustering were not considered. For case PA3, only a minority of nuclei had malignancy scores > 2 standard deviations above the mean. Therefore, these cells were identified using outlier detection in a normal distribution as done in the getOutliers(, method = “I”, rho=c(0.1,3))$iRight) in R. getOutliers is part of extremevalues R package https://github.com/markvanderloo/extremevalues. Only the consensus nuclei that were identified as CNV positive in both approaches were considered for analysis. Less than 7.0% of the nuclei were called alternately by the two methods and were excluded from the analysis. Identification of CNVpos nuclei in recurrent glioma samples was conducted through a combination of InferCNV and identification of clusters with high expression of tumor markers SOX2 and PTPRZ1.

### Survival Analysis

Survival analysis was performed using the survfit() function in the survival package in R [51, 52], using the binarized enrichment of each of the gene sets as the covariate in the formula. For cox proportional hazards, the function coxph() was used in R, and the covariates are indicated in the main figures.

### Correlation Analysis

Correlation analysis of glioma proportions was done using Pearson correlation (function cor() or psych::corr.test in R). Correlation heatmaps were generated using the corrplot package in R.

### Identification of glioma state and lineage top gene markers

The lineage specific genes were determined using scater::findmarkers(…, direction =”up”) function on the top-level lineages (Neurons, astrocytes, microglia, undetermined, oligodendrocytes, OPC, and endothelial cells). The glioma-state specific genes were determined using scater::findmarkers(…, direction =”up”) function on the neoplastic glioma states only. To select specific lineage/glioma state markers, we further filtered the top markers generated above by selecting the genes with positive log-fold change values in 90% or more of the cluster-to-cluster comparisons. The top 150 genes were selected and are provided in **Supplementary Tables 2-4** for primary glioma, recurrent glioma, and non-neoplastic lineages, respectively.

### Principal component analysis

PCA analysis was done in factominer R package[28]. A matrix of snRNAseq sample by cell type/cluster was used as input (**Supplementary table 1)**. The proportion of each snRNAseq sample with respect to 11 non-neoplastic cell types (Ast1, Ast2, Ast3, Endothelial, mgTAM, moTAM, Myel1, Neuron, Oligodendrocyte, OPC, prTAM, Tcell) and the summed proportion of all neoplastic cell types (CNVPos) were used to form the initial PCA axes. The proportions of CNVpos cells that were assigned to each of the glioma states (gl_Mes1, glMes2, gl_Pro1, gl_Pro2, gl_PN1, gl_PN2) were added as supplementary quantitative variables.

PCA coordinates of the bulk RNAseq validation dataset were generated using Factominer, by using the normalized expression of the genes filtered by rowsum>1000.

### Acquisition of Tissue and Preparation of Acute Slice Cultures

Primary GBM tissue from two separate surgeries, TB 6571 (3 blocks of tissue) and TB 6579 (2 blocks of tissue) (see supplemental table for related clinical information), performed at Columbia University Medical Center/New York Presbyterian Hospital were retrieved fresh from the operating room in a sterile specimen cup and transported back to the laboratory on ice. Primary GBM acute slice cultures were prepared exactly as described previously [53]. Slices were treated as described previously with either DMSO or 5µg/mL Cerulenin for 18 hours prior to preservation and RNA extraction.

### Bulk RNAseq using Plate-Seq

RNA extraction was done using the RNeasy Mini Kit (Qiagen cat# 74106). RNAseq was on spatially localized biopsies was performed using Plate-seq as described [54]. 75bp paired end sequencing was performed on Illumina NextSeq platform and read alignment was done using STAR [55] to the human genome (hg19, annotation: UCSC known genes), and analysis was done as previously described [3]. FPKM values were used in GSEA analysis. The count matrix for the TCGA GBM dataset was downloaded using the GDCquery tool in R. The Chinese Glioma Genome Atlas (CGGA) RNAseq datasets [56, 57] was downloaded from (http://www.cgga.org.cn/download.jsp). The counts were normalized using the vst() function in deseq2 R package [58]. IDH-WT only samples were kept from both datasets (TCGA: 139 samples, CGGA: 179 samples) and used for downstream analysis.

For acute slice-culture PLATE-seq analysis, slices were then transferred to OCT and frozen into blocks. Tissue from each slice was isolated for RNA extraction by the Columbia Molecular Pathology Core using QiaSymphony extraction method. Total RNA was quantified using Nanodrop measurements, and 150ng of RNA from each slice/condition was loaded into a well of a 96 well plate. Pooled library amplification for transcriptome expression (PLATE-Seq) was then performed on the 96 well plate as previously described [54]. FASTQ files were demultiplexed and aligned to reference genome and transcript counts were normalized via DESeq2. U87 and Astrocyte co-culture PLATE-Seq was conducted the same way as described above.

### Differential gene expression analysis

For comparing astrocytes to glioma, EdgeR glmQLFTest was used and the top 3000 differentially expressed genes with an FDR cutoff of 25% [59] were extracted. Only datapoints with adjusted p-values less than 0.05 were used in downstream analysis. For plate-seq data differential gene expression analysis between treatment and control was performed adjusting for tissue block and patient using the Deseq2 pipeline [58]. For astrocyte cultures, differential gene expression analysis between treatment and control was performed adjusting for astrocyte passage and cell culture batch using the Deseq2 pipeline.

### Geneset enrichment analysis and Gene Ontology Analysis

The average normalized counts per gene per cluster was calculated. The resultant cluster-wise count matrix was used as input to the GSVA pipeline [60]. Gene sets used for various tests are provided in the supplementary material (**Supplementary Table 2-4**). The options used for performing the GSVA pipeline are as follows: method= ssgsea, kcdf=“Gaussian”, mx.diff=TRUE. Heat maps were generated using the heatmap.2 in R function from the package gplots (R Package) and scores z-scaled were indicated. Ontology enrichment analysis in gProfiler with default settings [61]. For GSEA of the combined TCGA and CGGA dataset and the validation bulk RNAseq dataset (Figure 3C) and Figure 3I, the enrichment was performed using method = “gsva” option on normalized counts, which normalizes the enrichment scores for each gene set per sample. GSEA analysis in Figures 3G and 4F was conducted using pre-ranked GSEA and was performed as described in Subramanian et al 2005 with 1000 permutations [62]. Log-2-fold-change between cluster B and the remaining clusters was used to rank the genes for the analysis, and marker genes from each sub-lineage was used for the gene sets. For GSEA in Figure-4F, preranked GSEA (based on log2foldchange) was performed using tissue cluster gene sets and the genes significantly upregulated after astrocyte treatment with cerulenin or the top 150 genes unique to tissue state B signature.

### Generating the tissue state signatures

Pseudo-bulk samples from the snRNAseq dataset were created by summing and rounding the normalized counts per sample. Differential gene expression analysis using the DESseq2 pipeline was conducted between the clusters, controlling for sequencing batch (Supplementary table 3). The genes significantly differentially increased in cluster C vs A and C vs B constituted tissue state C signature. The genes significantly differentially increased in cluster B vs A and B vs C constituted tissue state B signature. The genes significantly differentially increased in cluster A vs C and A vs B constituted tissue state A signature. Clustering samples of the validation bulk RNAseq dataset into the three tissue states was performed by first retrieving the geneset enrichment scores of the genes unique to each tissue state in each sample using the gsva algorithm with method=”gsva”. Next, we performed hierarchical clustering on the Euclidian distance matrix calculated from the gsea scores, with method=”Ward.D2” in hclust, and cutree function with k=3 – all in R.

### Spatial transcriptomics

Spatial transcriptomics was conducted using 10X™ Visium Spatial Gene Expression Slide & Reagent Kit, 16 rxns (PN-1000184), according to the protocol detailed in document CG000239_RevD available in 10X demonstrated protocols. 10 micron-thick tissue sections were mounted on the ST slides and stained for nuclei – DAPI among other antigens using a rapid immunofluorescence protocol described in document CG000312_RevB available in 10X demonstrated protocols. Imaging of whole slides was done at 20X magnification on a Leica Aperio Versa scanner or a Leica DMI6 thunder tissue imager. After imaging, the slides were de-cover-slipped and the tissue permeabilized for 11 minutes (which was empirically determined to yield best results based on the Visium Spatial Tissue Optimization Slide & Reagent Kit PN-1000193 as detailed in the protocol provided in document CG000238_RevD available in 10X demonstrated protocols). The remaining steps were conducted according to the manufacturer’s protocol. The libraries were sequenced on multiple Illumina Nextseq 550 (paired end dual-indexed sequencing) flowcells to achieve the recommended number reads per ST spot. The spatial transcriptomic (ST) samples were prepared using 10X genomics Cell Ranger (version 6.1.2) and Space Ranger (version 1.2.1) software. Raw tiff images of the tissue were labeled with Cell Ranger which generated a json file for Space Ranger to use during alignment. Labeled spots from Cell Ranger were inputted into the loupe-alignment argument in Space Ranger along with its respective tiff image file, FASTQ reads, and slide numbers. The reference genome used for alignment was built using the Space Ranger function spaceranger mkgtf with GRCh38 as the assembly and Ensemble 91 for the transcript annotations. All other parameters to generating the counts data for ST were set to its default setting. The number of counts per spot per ST sample is shown in **Figure S10C**. The plots of ST experiments shown in **Figure 2D** and **Figure S11** were generated using SPATA2 [63].

### Deconvolution and Spatial Cross Correlation Analysis

Deconvolution using *RCTD* was used to determine the proportion of each cell type at each spot in each of the 9 ST experiments[29]. *RCTD* was run in “full” mode and used the complete annotated set of single nuclei (*n=*43,505) as a reference. The differential gene expression threshold in the “createRCTD” step was set to 1.25 logFC; other parameters were set to their default values. Bulk samples were deconvolved by supplying null coordinates as described in the package documentation. Deconvolution performance was quantified using immunohistochemical staining of the ST samples for DAPI **(Supplementary table 1)**. Using the package BayesSpace, spots in each experiment were clustered according to their transcriptional profiles and cartesian coordinates. The number of BayesSpace clusters in each sample was determined using the SC.MEB package with criterion set to “MBIC”[64]. The mean proportion of each cell type was calculated within each BayesSpace cluster (total clusters n=33). Using QuPath cell detection, cells were segmented and then assigned to each BayesSpace cluster using the st_within function (R package *sf*). The number of DAPI-positive nuclei within a BayesSpace cluster was computed and divided by the area of the BayesSpace cluster to obtain the nuclear density. The density was then correlated with the mean proportion of each cell type and CNVpos (the sum of all neoplastic cell types) using the cor.test function.

Cohabitation patterns between cell types in the ST data were quantified using spatial cross correlation as implemented in the R package *MERINGUE* and evaluated at neighborhood size = 900uM [65]. To determine the spatial adjacency matrix for the first order calculation (a spot’s immediate neighbors), the Cartesian coordinates of each Visium spot were input into the “getSpatialNeighbors” function. Because Visium spots are arranged in a hexagonal lattice, the parameter “filterDist” was initially chosen such that no spot had greater than six contiguous neighbors. Spatial adjacency matrices were created using the *igraph* package by generating a graph from the first order spatial adjacency matrix using the “graph_from_adjacency_matrix” function and then inputting this graph into the “connect” function [66].

The deconvolved proportions of each cell type output by *RCTD* (summed to unity on a spot-by-spot basis) and the first order spatial adjacency matrix were used as inputs for the “spatialCrossCorMatrix” function in *MERINGUE* to determine the average pairwise spatial cross correlations between each spot and its neighbors between each of the 18 cell types (171 combinations total) in each experiment, The pairwise comparisons were normalized with respect to the number of spots in each sample before being averaged across samples and plotted using the *ComplexHeatmap* package[67]. The significance of the cross correlations was determined using the “spatialCrossCorTest” function with 100 permutations: the spatial cross-correlation calculation was repeated 100 times using neighborhoods consisting of randomly selected spots from that sample. The proportion of random permutations that yielded spatial cross-correlations at least as high as those obtained from the actual data was taken as the p-value for that relationship. For each sample, adjusted p-values for each cross-correlation relationship were determined using Benjamini-Hochberg correction. Significance across samples was computed using the Fisher method for combining p-values across independent experiments (*poolr* package, R, unweighted). Relationships with a combined adjusted p-value less than 0.05 were considered significant. Dendrograms for each of the resultant heatmaps were determined using ward.D clustering of the Euclidean distance between the spatial-cross correlation values for each cell type.

### Immunohistochemistry, histology, and in situ hybridization

Standard chromogenic Immunohistochemistry was done as described previously [23]. Paraffin-embedded formalin-fixed tissue sections or fresh frozen sections briefly fixed in 4% PFA, for 10 min (40 C) in 4% PFA in PBS. Paraffin sections after deparaffinization were treated with antigen unmasking solution according manufacture recommendations (Vector Laboratories, Burlingame, CA). The following antibodies and dilutions were used SOX2 (1:200, Mouse monoclonal, Abcam, Ab218520), KI67 (1:500, rat monoclonal polyclonal, Thermo Scientific, 14-5698-80), CD68 (1:200, mouse monoclonal, Abcam cat# ab955), NeuN (1:1000, mouse monoclonal, Millipore, MAB377). For fluorescent IHC, secondary antibody conjugated to fluorophores: anti-mouse Alexa Fluor 488 and 594 and anti-rabbit Alexa Fluor 488 and 594; goat or donkey (1:300, ThermoFisher Scientific, Eugene, OR) were applied for 1 hr at room temperature. In situ hybridization was done using RNAscope™ multiplex fluorescent v2 (ACDbio cat# 323100) per the manufacturer’s protocol in 5-micron paraffin-embedded, formalin-fixed tissue sections. We used predesigned probes for PTPRZ1, CLU, TOP2A, NOVA1, MEG3, and SOX2 from ACDbio; cat# 584781, 584771, 470321, 400871, 584801, and 400871, respectively. Fluorescent images were taken on a Zeiss 810 Axio confocal microscope at 40X. Brightfield fluorescent images were taken on an Aperio LSM™ slide scanner at 20X and 40X.

### Quantification of ISH

For quantification of in situ hybridization images we used the positive cell detection function in Qupath v0.2.3 [68]. We only quantified signal contained in DAPI-positive nuclei. First, DAPI positive nuclei were detected using the cell detection tool. Next, subcellular detection function was employed to segment puncta per each of the three probe channels. A random tree classifier was used to classify nuclei to be positive or negative in QuPath under default settings, with a minimum of two puncta per channel to classify a nucleus as positive for the probe. Infiltrated cortex and cellular tumor core were annotated by a neuropathologist.

### Cell Culture and co-culture

Human Astrocytes (ScienCell cat #1800) were cultured in Astrocyte culture medium (ScienCell cat# 1801), 2% fetal bovine serum (ScienCell cat #0010), 1% astrocyte growth supplement (ScienCell cat# 1852) and 1% penicillin/streptomycin (ScienCell cat # 0503). The cells were maintained as adherent cultures on poly-L-Lysine coated tissue culture plates. The cells were passaged at 70-90% confluence and treated at passage numbers 5-7. DMSO or Cerulunin Sigma cat#C2389 at 5ug/ml was used to treat the cells for 18hours as indicated.

Human astrocytes used co-culture were first transduced with lentiviruses carrying GFP (LentiORF control particles of pLenti-C-mGFP-P2A-Puro Origene™ cat# PS100093V), CLU (Lenti ORF particles, CLU (mGFP-tagged)-Human Clusterin (CLU), transcript variant 1 Origene™ cat# RC211875L4V), or LGALS3 (Lenti ORF particles, LGALS3 (mGFP-tagged) – Human lectin, galactoside-binding, soluble, 3 (LGALS3), transcript variant 1, Origene™ cat# RC208785L4V). Transduction was performed by inoculating astrocytes seeded at 10^4 cells per well with 10 ul of virus at 1×10^7 TU/ml (5 MOI), in the presence of 10ug/ml polybrene, followed by one-week selection in a 0.5ug/ml puromycin containing selection medium. Transduction efficiency was confirmed by observing fluorescence on microscopy and FACS analysis. Co-culture with U87-MG (obtained from ATCC – maintained in DMEM+10%FBS) ensued for 24 hours – with both Astrocytes and U87-MG cells seeded at 2*10^5 cells/well – 6-well plate). The cultures were trypsinized 24hrs after seeding and subjected to FACSorting (influx cell sorter - Beckman Coulter, Jersey City, NJ), into GFP+ (astrocyte) and GFP- (U87MG) fractions, from which RNA was extracted as described above.

### Real time quantitative PCR

Total RNA was extracted from brain specimens using RNAeasy minikit (Qiagene©). RNA concentration and purity were determined using NanoDrop (Thermo Scientific™, MA). RNA was converted to cDNA using High capacity RNA-to-cDNA kit (Thermo Fisher Scientific, Applied Biosystems™, MA). The following Taqman assays were used (CHI3L1 - Hs01072228_m1, CD44 - Hs01075864_m1, LGALS3 - Hs00173587_m1, GAPDH - Hs02786624_g1, MIB1 - Hs01075903_m1, HES5 - Hs01387463_g1, CLU - Hs00156548_m1, EZH2 - Hs00544830_m1, SOX2 - Hs04234836_s1, NES - Hs04187831_g1, HES1 - Hs00172878_m1). The reaction volumes were 15 ul per reaction. TaqMan™ Multiplex Master Mix (Thermo Fisher Scientific cat# 4461881) was used. All reaction included 5ng of cDNA. Thermal cycling parameters were conducted per manufacturer’s standard recommendations. The qPCR plates were read on a QuantStudio™ 5 Real-time PCR system (Thermo Fisher Scientific, Applied Biosystems™, MA). The reactions were done in triplicates. Relative gene expression was calculated using the delta delta Ct method with GAPDH as a reference gene.

### Statistical testing

Statistical comparisons were done using one-way ANOVA (or Kruskal Wallis test) and Tuckey post-hoc comparison in R. Statistical testing for RNAseq application is reported in the main text or respective methods section. Differential abundance analysis was done employing a moderated regression model in ANCOMBC with default parameters, assigned Condition (primary vs post-treatment recurrence) and CNVpos proportions in the design formula, and as described by the authors ([27]). One tailed paired t-tests were done to compare the core and margin percentages of the same case (Figure 3). A one sample t-test was conducted to determine if the percentage of TOP2A+ that were CLU+ was less than 50%. One or two tailed t-tests were used in Figure S13 as indicated in the legend.

### Data availability

Data for spatial transcriptomic can be queried using an interactive web app: https://vmenon.shinyapps.io/gbm_expression/. All other data will be deposited as in GEO repositories prior to publication.

## Supporting information

Supplementary_table_1_metadata_composition_ST_diagnostics_cell_staining

Supplementary_table_2_primary_glioma_markers_all_clusters_GO

Supplementary_table_3_rec_glioma_markers_all_clusters_GO

Supplementary_table_4_Astrocytes_markers_all_clusters_cnvneg_GO_subcluster_2

Supplementary_table_5_myeloid_markers_combined_clusters_cnvneg_GO

Supplementary_Table_6_CERULENIN treatment_astrocytes_slicesCx

Supplementary_table_7_snRNAseq_comp_Clusters_signature_genes

supplementary_table_8_in_vitro_astrocytes_CLU

## Acknowledgment

The results published here are in part based upon data generated by the TCGA Research Network: https://www.cancer.gov/tcga. This research was funded by NIH/NINDS R01NS103473 (PC, PAS, JNB), The William Rhodes and Louise Tilzer-Rhodes Center for Glioblastoma at New York-Presbyterian Hospital (OA, VM, PC), Herbert Irving Comprehensive Cancer Center pilot research grant (OA), and the NIH/NCI Cancer Center Support Grant P30CA013696 (OA, ABL, JNB, PAS, PC).

This research was funded by NIH/NINDS R01NS103473 (PC, PAS, JNB), William Rhodes and Louise Tilzer-Rhodes Center for Glioblastoma (RCG) Collaborative Research Initiative grant (OA, VM, PC), HICCC pilot research grant (OA), and in part through the NIH/NCI Cancer Center Support Grant P30CA013696. This research was supported by the Genomics and High Throughput Screening Shared Resource and the Digital Computational Pathology Laboratory in the Department of Pathology and Cell Biology at Columbia University Irving Medical Center. We thank Dr. Claudia Deoge for help with RNAscope. We thank the immunohistochemistry core in the Department of Pathology and Cell Biology at Columbia University Irving Medical Center for help with GFAP-SOX2 immunostains. We thank Al-Fardthakh© for support with aspects of bulk RNAseq data analysis.

## Author contribution

These authors contributed equally: Osama Al Dalahmah, Michael Argenziano, Adithya Kannan

Corresponding Authors: Osama Al-Dalahmah, Vilas Menon, Peter Canoll

OA, JNB, PAS, VM, and PC designed the study; OA, MA, AK, DB, AS, AM, FK, JFT, ARG, AT, MAB, JF, TS, EB, JNB and PC conducted the experiments, including tissue procurement, tissue processing, and tissue analysis. OA, MA, AK, PAS, VM, and PC analyzed the data. OA, MA, AK, VM, and PC wrote the manuscript and all authors edited, read, and approved the final manuscript.

## Supplementary Results

### Single nucleus RNAseq reveals proliferative, astrocyte-like/mesenchymal, and progenitor-like/proneural states in both primary and recurrent GBM

Radiographically, GBM typically has a CE core surrounded by a non-enhancing infiltrated brain that is highlighted by FLAIR-signal abnormality by MRI (**Figure-S1A**). The histopathological features of the resected tumor can vary from highly cellular tumor with vascular proliferation to less cellular infiltrated brain. These features are shown in **Figure-S1H**, demonstrating samples with a cellular GBM core (red star in **Figure-1A, Figure-S2H PA1, PA2, PA3, and PO2_1**) and others with overlying cortex (green star in **Figure-S1A, Figure-S2H PO2_2 and PO1**), which we use below.

To explore the heterogeneity of primary GBM, we analyzed several banked surgical samples using snRNAseq as shown in (**Figure-S1A**). A total of 8 samples from 7 patients were selected for analysis (**Supplementary Table-1**). Neuropathological assessment of tumor cellularity ranged from cellular tumor with hallmarks of GBM, to reactive brain parenchyma with few atypical cells. This assessment was made on Hematoxylin and Eosin (H&E) stained formalin fixed paraffin embedded sections adjacent to or frozen cryosections of the frozen tissue analyzed by snRNAseq (**Figure-S2H)**. We isolated nuclei from the frozen tissue and subjected them to snRNAseq followed by downstream analyses including clustering, differential gene expression analysis, cluster marker detection, and gene set enrichment analysis (GSEA) as outlined (**Figure-S1A**). 15189 nuclei passed our QC (**Supplementary Table-1**). To distinguish putative glioma cells from non-neoplastic cells, we employed an established approach that infers large scale copy number alterations/variations (CNV) from RNA expression profiles [49]. Chromosomal heat maps showing putative neoplastic nuclei are shown in **Supplementary Figure-S2A-G**. Next, we also applied a second method to label nuclei based on a “malignancy score”, which we have previously shown to be a robust metric to distinguish glioma cells from non-neoplastic cells [2, 10], and the consensus nuclei designated by both methods was used for downstream analysis. Nuclei with no consensus CNV status were excluded (4.7%). Uniform manifold approximation and projection (UMAP) plots from individual cases labeled by transformation status are shown in **Figure-S1B**. We identified 7954 putative neoplastic nuclei with inferred large scale chromosomal CNV (CNVpos/glioma nuclei). Glioma nuclei showed multiple chromosomal alterations including gains of chromosome 7 and losses of chromosome 10 (**Figure-S1**). Having identified neoplastic and non-neoplastic nuclei, we aligned the datasets from multiple samples and performed clustering analyses separately on CNVpos (glioma) nuclei from all cases using shared nearest neighbor and the smart local moving algorithm [69]. A UMAP plot is shown for all primary glioma nuclei non-neoplastic nuclei color-coded by glioma state/lineage **Figure-S1C**. This approach identified 6 distinct clusters: these resembled progenitors (oligodendrocyte-progenitors (gl_PN1 - proneural) and neural-progenitors (gl_PN2 - proneural), astrocytes (gl_Mes1 and gl_Mes2 - mesenchymal), and proliferative cells (gl_Pro1 and gl_Pro2).

The identity of the glioma states is akin to previously described glioma states, as demonstrated by the enrichment of several gene lists from [3, 9, 70, 71] **– (Figure-S1D, supplementary Table-2)**. For example, gl_Pro1 and gl_Pro2 showed enrichment in gene sets specific for cell-cycle phases [71], with gl_Pro1 showing highest enrichment of G2/M genes (Gobin_G1) and gl_Pro2 showing highest enrichment of G1/S phase genes and DNA repair related genes (Gobin_G3). Clusters gl_PN1 showed enrichment of the Verhaak’s proneural, and OPC signature genes, while gl_PN2 showed enrichment of NPC signature genes. Finally, gl_Mes1 showed enrichment of astrocyte-like signatures and Verhaak’s classical signature while cluster gl_Mes2 showed enrichment of several gene sets related to reactive astrocytes, and Verhaak’s mesenchymal signature [3, 7, 70]. Our clustering is consistent with that described in Neftel et al. 2019 [9] and Wang et al. 2019 [7], and the states we describe are compatible with those in Yuan et al 2018 [2]. To further clarify the cellular phenotypes represented in our glioma clusters, we measured the enrichment of the major biologic process and molecular function gene ontologies (GO) in the glioma state top gene markers (see methods). GO enrichment analysis demonstrated enrichment of GO’s relating to locomotion, neurogenesis, neuronal migration, and cell projection in gl_PN1 markers genes; Notch signaling, neuron development, and GABA reuptake differentiation, and synaptic signaling in gl_PN2 genes; response to organic substances, ion homeostasis, and Signaling by tyrosine kinases in gl_Mes1 genes; response to cytokines, interferon gamma, and leukocyte activation and immune response in gl_Mes2 genes; mitosis and nuclear division in gl_Pro1, and S-phase, DNA replication, and DNA repair in gl_Pro2(**Figure-S1F** and **Supplementary Table-2**). The identities of the clusters can also be appreciated by examining select gene markers **Figure-S1E and Supplementary Table-2.** gl_Pro1 expressed cell-cycle genes TOP2A, CENPF, and AURKB. gl_Pro2 showed highest expression of DNA damage/repair including FANCI, HELLS, and XRCC2. gl_PN2 showed high levels of CD24, MEG3, and SOX4. gl_Mes1 showed high levels of protoplasmic astrocyte genes including SLC1A3, LIFR, ATP1A2, C1orf61, and NTM, while gl_Mes2 showed highest expression levels for reactive astrocyte and immune genes including CLU, VIM, and SAT1. While our glioma states resemble those described in the literature, less is known about whether glioma cells assume similar states in the recurrent setting. Therefore, we bridged this gap by directly analyzing recurrent IDH-WT glioma samples using the same approach we used for primary GBM samples.

To define the states of IDH-WT glioma in the post-treatment recurrence setting, we analyzed 8 cases of post-recurrent IDH-WT glioma using snRNAseq (**Figure-S3A**). We identified 8908 neoplastic nuclei harboring large-scale CNV (**Supplementary Figure-S4**). Of the eight cases, two were paired recurrences from the primary samples (TB5124 – recurrent of TB4916, and TB5053 – recurrent of TB4718, see respective section on comparing paired samples below). We treated recurrent gliomas similarly to the treatment naïve primary tumors and clustered all neoplastic nuclei together. Like primary gliomas, we found that recurrent glioma clusters can be assigned two proneural, two mesenchymal, and two proliferative states (**Figure-S3B**). The gene markers of the recurrent glioma states are enriched for similar ontologies to those seen for primary glioma states (**Figure-S3C** and **Supplementary Table-3**), showed similar patterns of enrichment for the previously presented gene sets in **Figure-S1D** (**Figure-S3D**), and displayed comparable gene marker expression (**Figure-S3F**). These results demonstrate that post-treatment recurrent glioma states closely resemble states observed in the primary pre-treatment setting. Indeed, Pearson correlation analysis demonstrates that corresponding states were positively correlated (**Figure-S3E**). The correlation patterns reveal that gl_Mes1 and gl_Mes2 are positively correlated with each other in the primary and recurrent settings. This is also seen with gl_PN1 and gl_PN2, as well as gl_Pro1 and gl_Pro2. We therefore contend that a view of primary and recurrent glioma states may benefit from simplification and embrace a viewpoint that primary and recurrent glioma states can be classified as progenitor-like/proneural (gl_PN1 and gl_PN2), astrocyte-like/mesenchymal (gl_Mes1 and gl_Mes2), and proliferative (gl_Pro2 and gl_Pro1) states. A select set of markers of both primary and recurrent GBM states is provided in **Figure S4I**. Assigning cell cycle scores using Seurat cell-cycle score assignment reveals that gl_Pro1 has the majority of cells in G2M phase, whilst gl_Pro2 has the majority of cells in S phase **Figure S4I**. Integration of both primary and recurrent glioma nuclei shows cells from primary and recurrent samples overlap in the UMAP space, and that this overlap is seen for all 6 GBM states (**Figure S5G**).

While the transcriptional signatures of glioma are relatively well defined, the spatial distribution of these glioma states is less well understood. Given the marked difference in cellular composition between the cortex and the deeper (typically more heavily infiltrated) white matter, and the highly cellular tumor core, we asked if these different anatomic regions harbor distinct glioma states. In other words, we posited that the cellular microenvironment of glioma influences glioma states. Specifically, we hypothesized that we would find more glioma cells that resemble astrocytes (astrocyte-like/mesenchymal glioma) or neurons (progenitor-like - specifically gl_PN2) in the cortical margins. To address this question, we examined the expression of select combinations of glioma state transcripts using in situ hybridization (ISH) across the cellular tumor and the infiltrated cortical margin. We used probes to detect PTPRZ1 (high in glioma), CLU (high in astrocytes and astrocyte-like/mesenchymal glioma), SOX2 (high glioma), NOVA1 (high in progenitor-like/proneural glioma), and MEG3 (high in neurons and progenitor-like/proneural glioma - gl_PN2) in the cellular core and overlying infiltrated cortical margin in 5 cases of primary GBM (**Figure-S5A, C**). We found that significantly higher proportion of PTPRZ1+ glioma cells co-expressed CLU in the cortex versus the core (**Figure-S5B**). Similarly, we found that significantly higher proportion of SOX2+NOVA1+MEG3+ glioma cells in the cortex versus the core (**Figure-3D**). These findings indicate that the different glioma states have distinct distributions throughout the landscape of glioma and suggest that local tissue cellular composition and perhaps other microenvironmental influences can affect glioma states. We note that astrocyte-like/mesenchymal glioma states were negatively correlated with proliferative states. Consistent with this result, our ISH findings demonstrated a significantly smaller proportion of CLU+ cells that co-expressed TOP2A (mean=31.71388837%, Standard deviation = 15.73850618, one-tailed t-test p= 0.000249641, n=5, **Figure S5E-F**).

### Comparison between primary and recurrent glioma pairs

Not surprisingly, the recurrent tumors did not show identical chromosomal CNVs with their primary counterparts. While TB5014 retained the CNV of the TB4916 (gain of 7, loss of 10 and 14) and acquired additional alterations including gains in chromosomes 19 and 20 (**Figures S2F and S4A)**, TB5053 showed a complex gains and losses across multiple chromosomes (**Figures S2G and S4B)**.

In the main text, we note that gl_PN1 is depleted from our recurrent GBM samples (**Figure 2A**). This is consistent with the literature [6], since the Verhaak classical subtype resembles our gl_PN1, which showed positive enrichment scores of the Verhaak’s classical gene set. Of the non-neoplastic cell types, OPCs were depleted in recurrence. This may be explained by the fact the OPCs are the proliferative cell type in the brain and glioma treatment with chemotherapy and radiotherapy depletes proliferative cells, as have been previously demonstrated [72].

### Analysis of low-grade glioma and epilepsy cases

To sample states of myeloid cells and astrocytes across different disease states, we chose to analyze the microenvironment of low-grade glioma (LGG) and temporal lobe epilepsy. We conducted snRNAseq on 6 cases: two IDH-mutant oligodendroglioma (TB3652 & TB3926), one IDH-mutant astrocytoma (TB4100), and three temporal lobe epilepsies (TB4189, TB4437, & TB4957). We identified 970, 1154, 1036 nuclei for LGG cases TB3652, TB3926, and TB4100, respectively. We identified CNVpos nuclei using a combination of chromosomal CNV, clustering, and tumor marker expression as shown in **Figure S6**. Cases TB3652 and TB3926 had typical chromosome 1p and 19q codeletions (**Figure S6A, D**), and harbored 817 and 942 CNVpos nuclei, respectively (**Figure S6B, E**). The tumor nuclei expressed tumor markers SOX2, EGFR, and PTPRZ1, and/or OPC markers DSCAM and TNR; myeloid cells expressed a CD74, C3, ITGAX (CD11c), ITM2B, and/or HLA-B; while oligodendrocytes expressed MBP and MOG (**Figure S6C, F**). 382 CNVpos nuclei were found in case TB4100, which did not harbor CNVs across most cells, and CNVpos nuclei were identified by clustering and marker expression as noted above. Of the epilepsy cases, we identified 2558, 179, and 138 nuclei in cases TB4189, TB4437, and TB4957, respectively. **Figure S7A-C** show marker expression in cases TB4437, TB4189, and TB4957, where markers of astrocytes (GFAP, AQP4, SLC1A2, SLC1A3), neurons (RBFOX3, MEG3, GAD1, and SLC17A6), myeloid cells (CD74, ITGAX, C3, ITM2B), oligodendrocytes (MBP, MOG, OPALIN, and CNP), and OPCs (DSCAM, TNR, SOX2, and PDGFRA). The CNVneg nuclei from all LGG and epilepsy cases were combined with those from primary and recurrent IDH-WT GBM and were analyzed as presented in the section below (myeloid cells) and main text (astrocytes).

### Astrocytes cluster into three distinct astrocyte cell states

Based on the resemblance to known astrocyte phenotypes we curated three gene sets (**Supplementary Table-4**), which represent three major astrocyte states (protoplasmic, reactive-1, and reactive-2), and then clustered astrocyte nuclei using Ward D2 hierarchical clustering on the Manhattan distance of the enrichment scores (overlaid on the 3D tSNE plots in **Figure1**), into a protoplasmic cluster (Ast1), and two reactive clusters (Ast2 and Ast3 – as described in the main text (**Figure 1C**). We asked whether our method of clustering astrocytes, described in figure 1, can result in similar clusters to more “unbiased” methods. Thus, we performed Louvain clustering on shared nearest neighbor graphs (created through igraph – k=500 – **Fig. S13A**). Examination of marker expression for each cluster demonstrate that Louv 2 is similar to Ast1 - with expression of protoplasmic genes, Louv 3 is similar to Ast2 – with expression of PLP1 and ribosomal genes, and Louv 1 is similar to Ast3 – with expression of C3 and CD44 (**Fig. S13B**). Examination of the overlap of astrocyte calls between the method employed in figure 1 and the “unbiased” clustering reveals that the unbiased Louv clusters overlap to high extent with those described in Figure 1, as described above (**Fig. S13C**). Therefore, we conclude that our clustering approach we employed in figure 1 is highly analogous to unbiased clustering.

### Analysis of myeloid cell states

Myeloid cells have been implicated in modulating glioma migration, infiltration, and progression [73]. We identified 5925 nuclei we classified as myeloid cells. Unbiased clustering revealed 8 subclusters which we then used to assign the specific myeloid lineages. We merged clusters with similar enrichment scores of gene sets representing microglia-derived tumor-associated macrophages (mgTAM), monocyte-derived TAMs (moTAM), proliferative TAMs (prTAMs), and T-cells as described in [15] - Myeloid subclusters 0 and 9 were combined as Myel1 (baseline), subclusters 1 and 3 – moTAM (monocyte derived TAMs); subclusters 2, 4, 6, and 8 as mgTAM (microglia-derived TAMs); subcluster 5 was kept as prTAM (proliferative TAMs); and subcluster 7 was kept as T-cells. The enrichment of these gene sets in the final myeloid states is provided in **Figure SE**. A subset of myeloid cells showed mixed enrichment scores across mgTAM, moTAM, and dendritic cells, and were considered baseline (referred to as Myel1). Overall, we classified 2678, 1346, 1364, 360, and 177 nuclei as Myel1, moTAM, mgTAM, prTAM, and T-cells, respectively, and these are shown in 3D tSNE space in **Figure S9A**. Myel1 state showed higher expression of SAT1, CEBPD, and GLUL (**Figure S9C**-top row). moTAM showed highest expression of CD163, MS4A4E, NHSL1, FMN1, and MSR1 (**Figure S9C**-2nd row). mgTAM showed highest expression of SORL1, RIN3, ITGAX, HS3ST4, and FRMD4A (**Figure S9C**-3rd row). prTAMs showed highest expression of CST3, MEF2A, DBI, PLXDC2, and DOCK4 (**Figure S9C**-4th row). Finally, T-cells showed highest expression of CD2, CD247, CD96, FYN, and SKAP1 (**Figure S9C**-5th row). Different myeloid states were accounted for different conditions (**Figure S9B).** While Myel1 was present in Epilepsy, primary and recurrent GBM, mgTAM was the main state found in LGG, but was also in primary and recurrent GBM. moTAM, T-cells, and prTAM were found in primarily in recurrent GBM (**Figure S9D)**. The gene-wise DGE between myeloid states and the myeloid state markers are provided in **Supplementary Table-5**.

### snRNAseq of samples used for ST – a validation dataset

Single nuclei from each ST patient were analyzed when available (n = 7). The nuclei were obtained, cleaned, and analyzed as described elsewhere. CNVpos nuclei were identified using inferCNV (**Figure S8A-G**, sample QC, number of nuclei per sample as well as lineage assignment is provided in **Supplementary Table-1).** They were classified as was done for the previously presented datasets. The CNVneg nuclei were then classified into cell types using the *singleR* package (de.method=”wilcox”) with the previously annotated single nuclei data set as a reference[74]. Proportions of nuclei per cell type are included in **Supplemental Table-1.** Using the compositional matrix of these samples, they were able to be classified into tissue states using k-means clustering with the centers of the discovery data set samples supplied as centers (**Figure S8I**). The integrated CNVneg nuclei are shown in a UMAP (**Figure S8J**).

### Spatial cross-correlation analysis of deconvolved cell type proportions

Our analysis of cell type composition in snRNAseq samples highlights prognostically relevant compositional patterns. To examine these patterns with spatial resolution, we analyzed 9 IDH-WT GBM samples using spatial transcriptomics and deconvolution (see methods). To validate the accuracy of our deconvolution, we compared the distribution of deconvolved cell type proportions to fluorescent staining for select cell type markers. **Figure S11A-B** show the deconvolved proportion of neurons in a subset of ST samples alongside fluorescent staining of the same samples for NeuN, a canonical marker for neurons – see **Figure S10B**. This highlights that our deconvolution approach was able to reflect patterns of spatial heterogeneity that were also suggested by fluorescent staining. **Figure S11C-D** shows deconvolved proportions of select cell types in 2 ST samples and shows that cell types whose proportions covary in each tissue-state show similar patterns of spatial heterogeneity to each other across multiple samples. We quantified and aggregated trends across our 9 ST samples using spatial cross-correlation and tested them for significance—see methods for details. To determine the relative representation of tumor within each ST sample and confirm the ability of our deconvolution approach to identify tumor, we correlated nuclear density (cellularity) obtained from immunohistochemical staining for DAPI (**Figure S10A**) with our deconvolved cell type proportions. BayesSpace was used to segment each sample into clusters containing transcriptionally similar spots (see methods). A total of 33 clusters were generated (**Figure S10D)**. The density of nuclei was obtained for all of these clusters across the data set, and we calculated the correlation with the deconvolved proportion of each cell type (**Supplementary table 1**). The total proportion of CNVpos cell types was positively and significantly correlated with density of nuclei (correlation: 0.388, p=0.025). **Figure S11E** shows a representative sample with DAPI staining and **Figure S11F** shows the same sample segmented by BayesSpace generated clusters and colored by the proportion of deconvolved CNVpos cell types present in that cluster.

### The spatial landscape of glioma associated tissue-states in primary and recurrent GBM

To understand the spatial landscape of primary and recurrent glioma, we mapped the distribution of our “tissue-state” signatures in space in primary and recurrent GBM. First, we tested one of our cases that we utilized for snRNAseq (PO2) and took 48 localized biopsies that we analyzed using plate-seq [54]. Immunofluorescence of frozen sections taken prior to analysis revealed a cellular DAPI-dense glioma core and a NeuN rich cortical margin (**Figure S12A**). We conducted GSEA analysis of our tissue-state signatures in the RNAseq data from the localized biopsies and mapped that against the location of the biopsies (**Figure S12B**). Tissue-state C signature was highest in the core, compared to tissue-state A signature, which was highest in the cortical margin. Tissue-state B signature showed a more patchy distribution with foci of enrichment in both the core and margin. Interestingly, the intermediate region between the core and cortex, showed mixed enrichment across all three tissue-states. This data highlights the anatomic localization of tissue-state signatures and underscores the heterogeneous patterns in the intermediate non-cortical “margin” region.

Next we performed deconvolution on a previously published dataset of bulk RNA sequencing from MRI-localized biopsies of primary and recurrent GBM, and control brain samples [3] to assess the abundance of neoplastic and non-neoplastic cell types in different radiographic regions of primary and recurrent GBM. Our results showed that in contrast enhancing regions of primary GBM the cell types associated with tissue states B and C were more abundant than cell types associated with tissue state A, while in the contrast enhancing regions of recurrent GBM the cell types associated with tissue state B were more abundant than cell types associated with tissue state C or A. The FLAIR+ samples in primary GBM showed a mixture of neoplastic and non-neoplastic cell types from all three tissue states, while the FLAIR+ samples from recurrent GBM showed predominantly non-neoplastic cells, with highest abundance of cell types of tissue state A. As expected, control samples were also predominately composed of non-neoplastic cell types associated with tissue state A. (**Figure S12C)**. We also assessed for the expression of tissue-state signature genes in these same samples. This analysis showed similar patterns to those of the deconvolved cell types.(**Figure S12C**). In summary, contrast enhancing regions in both primary and recurrent tumors predominantly represent neoplastic and reactive cell types, but the distribution of specific glioma subtypes varies between primary and recurrent tissue. Non-enhancing margins of recurrent GBM samples predominantly represent reactive/gliotic brain tissue with relatively low levels of tumor infiltration, whereas the non-enhancing margins of primary GBM can contain a wider range of pathological features, including regions of abundant glioma infiltration.

### Astrocyte CLU alters U87 glioma cell gene expression

In examining the cellular milieu co-inhabiting tissue state B, we focused on Ast3, an astrocytic state with high expression of Clusterin (CLU). Astrocytic CLU is known to reduce amyloid accumulation in mouse models of Alzheimer’s disease, and is thought to be neuroprotective[21, 37, 75]. CLU is upregulated in GBM and can protect GBM cells from radiation-induced apoptosis [76]. However, little is known about the interaction between CLU+ astrocytes (i.e. Ast3) and GBM. We first identified the genes that were significantly correlated with CLU expression (using the psych::corr.test R function) in astrocytes that have high CLU expression, defined as in the 3^rd^ and 4^th^ quantiles of normalized CLU levels. These include ATP1B2, F3, AQP4, GJA1, CHI3L2, CHI3L1, LGALS1, and LGALS3 among others (**Figure S13D** and **supplementary table-8**). Analysis of pathways enriched in CLU-correlated genes reveals they encompass Reactome and KEGG pathways involved in signal transduction, Rho GTPases, Hippo signaling, and translation (**Figure S13E**). With a testable Ast3 (CLU-high) astrocytic signature at hand, we modeled an Ast3-like astrocyte state *in vitro* by overexpressing CLU in human astrocytes (**Figure S13F**). As a separate experimental condition we overexpressed LGALS3. rt-qPCR analysis shows that merely co-culturing astrocytes with U87 glioma cells leads to reduction of astrocytic SOX2, NES, CLU, and HES5 expression. Rt-qPCR confirms CLU and LGALS3 overexpression in CLU- and LGALS3-astrocytes, respectively, and reveals CLU astrocytes increase HES5 expression, whereas LGALS3 increase NES expression, both when compared to GFP control astrocyte in the setting of U87 co-culture (**Fig. S13I**). CHI3L1, and Ast3 gene, was increased in both LGALS3+ and CLU+ astrocytes. Since both CLU+ and LGALS3+ astrocytes model some aspects of Ast3, and only CLU+ astrocytes significantly increased astrocytic CLU, we chose to use the CLU+ astrocytes as an Ast3-like model and analyzed those cells further. Comparing the genes differentially expressed between sorted CLU-overexpressing versus control astrocytes revealed 274 differentially expressed genes, including many that are positively correlated with CLU levels as defined by human glioma-associated astrocytes (**Fig. S13G** and **supplementary table 8**). These genes are enriched in KEGG/Reactome pathways that encompass Hippo signaling, and extracellular matrix organization (**Fig. S13H**). These results provide support for the resemblance between Ast3 cells and CLU+ astrocytes. Next, we focused on glioma cells and asked if co-culture of U87 glioma cells with astrocytes leads to altered glioma gene expression. rt-qPCR of sorted U87 glioma showed that merely co-culturing U87 glioma with astrocytes leads to increased SOX2 and HES1 expression. When co-cultured with CLU astrocytes HES5 is increased in U87 cells, whereas HES1 is reduced in U87 cells co-cultured with LGALS3 astrocytes (**Fig. S13J)**. RNAseq of U87 glioma co-cultured with control (GFP) astrocytes leads to enrichment of gene ontologies involved in monocyte differentiation and leukocyte migration (**Fig. S13K and supplementary table 8.** When co-cultured with CLU-astrocytes, the transcriptome of U87 glioma cells is enriched in ontologies involved in glial differentiation, neural precursor proliferation, and biosynthesis of unsaturated fatty acids (**Fig. S13L and supplementary table 8).** Together, these results show that astrocytes can exert different effects on glioma gene expression, and Ast3-like astrocytes promote a signature related to glial differentiation and precursor proliferation.

---

**Figure S1:**
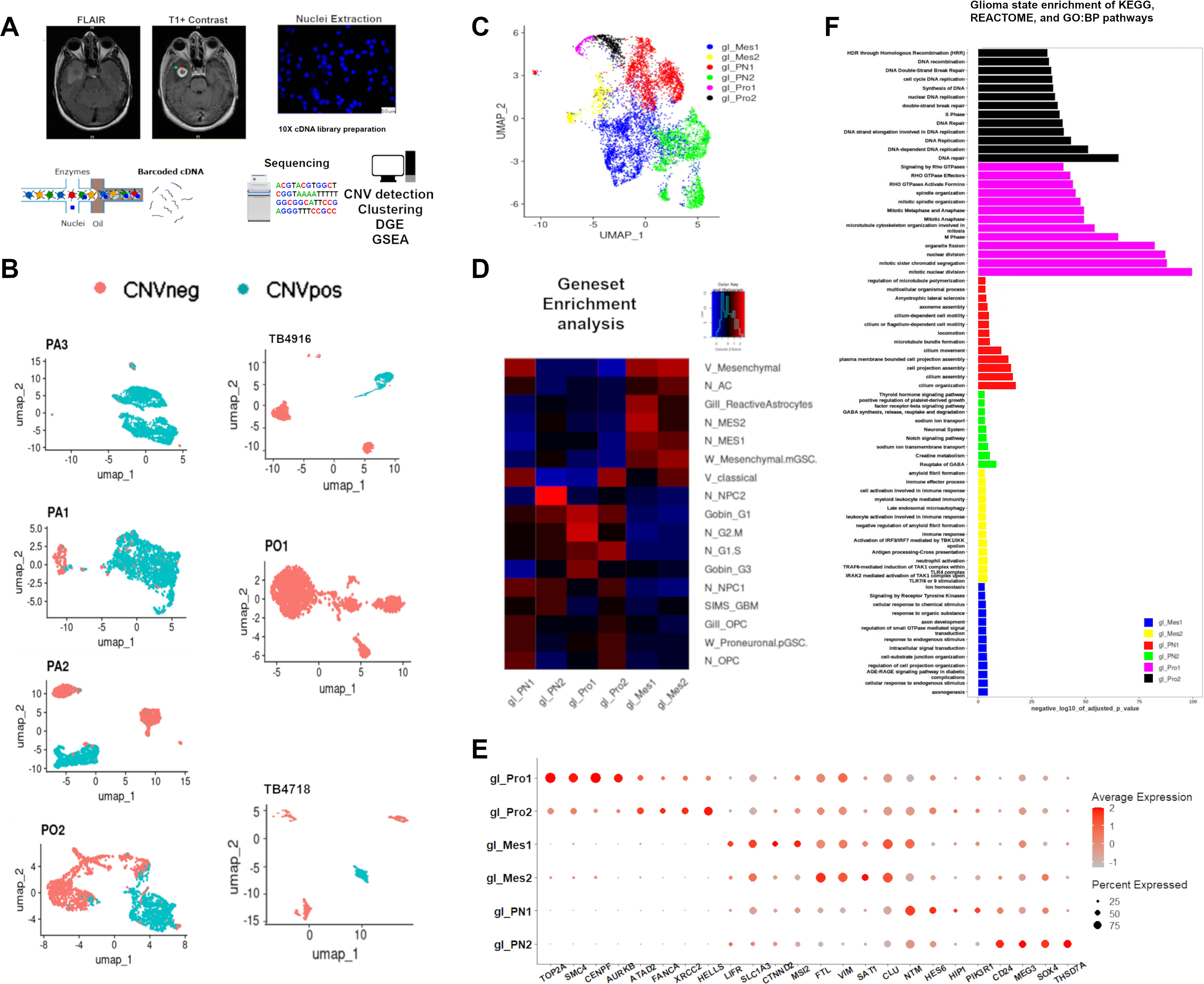
snRNAseq-derived transcriptional states of putative neoplastic nuclei from primary IDH-wildtype GBM samples. **A)** Outline of Analytic Design: T 2/FLAIR and post-contrast T1 MRI sequences of a glioblastoma showing the classic radiological appearance of a glioblastoma (Case PO2); with a ring enhancing mass (red star) with surrounding increased FLAIR signal (green star). The tumor was resected and banked (frozen). Nuclei are extracted from frozen tissue and are subjected to droplet based single nuclei RNA sequencing using the 10X chromium platform. The resultant barcoded cDNA is then sequenced and analyzed. Analyses performed include identification of putative neoplastic cells by identifying cells with inferred copy number variations (CNV), clustering, differential gene expression (DGE), and gene set enrichment analysis (GSEA). Scale bars = 50 um. **B)** Uniform-manifold approximation and projection (UMAP) graphs showing putative neoplastic (CNVpos) and non-neoplastic (CNVneg) nuclei from the seven primary IDH-wildtype glioma cases selected for analysis indicated by subpanels b1-b7. **C**) UMAP plot showing all putative CNVpos (**C**) nuclei from the seven primary glioma cases aligned and projected in shared UMAP spaces. The nuclei are color-coded by glioma state: Oligodendrocyte-progenitor-like (proneural - gl_PN1), Neural-progenitor-like (proneural - gl_PN2), Mesenchymal/astrocyte like (gl_Mes1 and gl_Mes2), and proliferative (gl_Pro1 & gl_Pro2). **D**) Geneset enrichment analysis (GSEA) of selected genesets from Verhaak et al. 2009 (v), Gobin M et al 2019, Gill et al 2014, Wang et al. 2019 (W), and Neftel et al. 2019 (N) showing enrichment of genes specific for states described in the literature in our described glioma states. **E**) Gene ontology (GO) term enrichment analysis (KEGG and REACTOME pathways and biological process GO) of the major terms enriched in glioma state top gene markers. The bars represent the negative log10 of the false discovery rate adjusted p.value, and are color-coded as in **C**.

**Figure S2:**
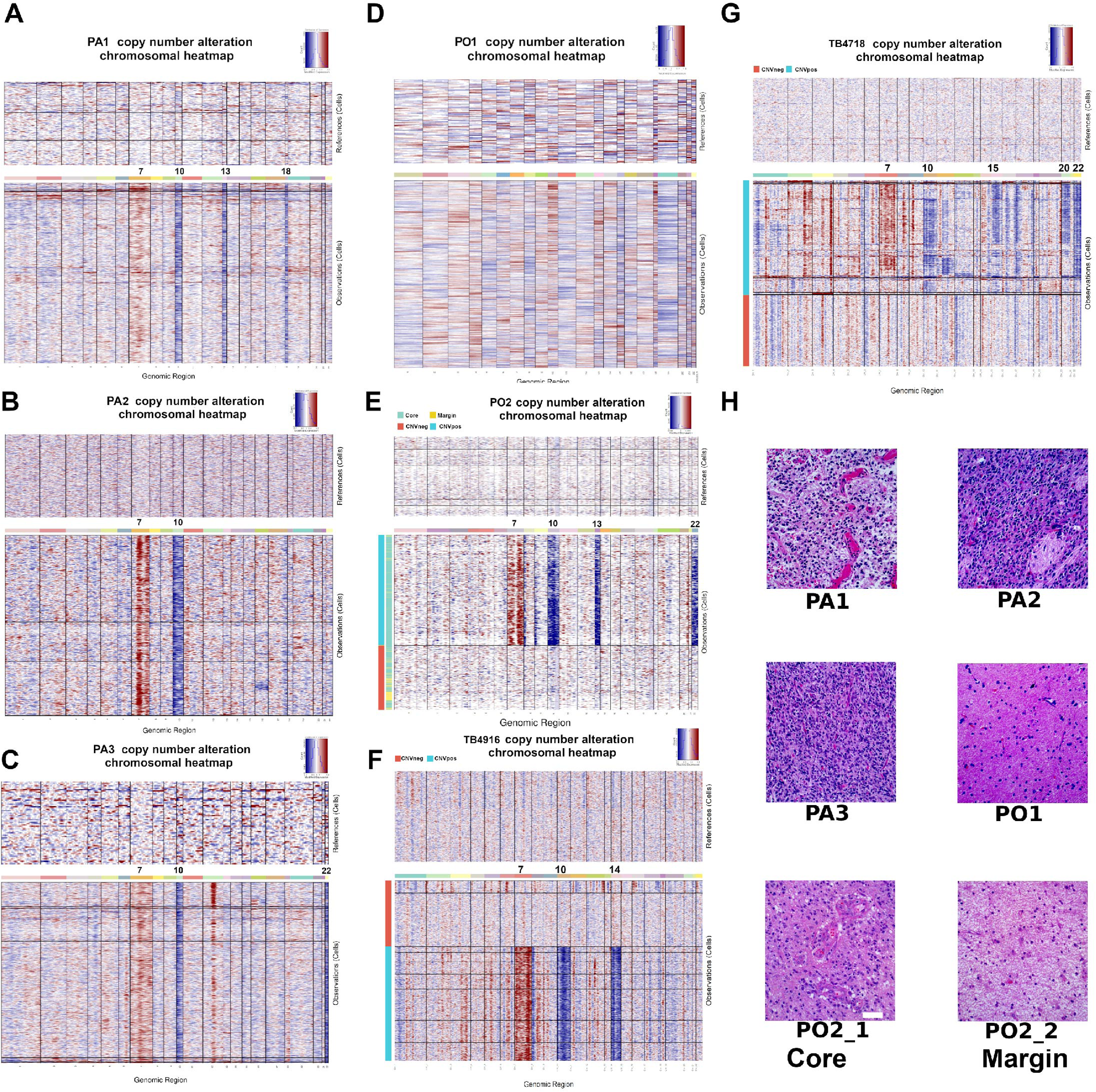
Identifying neoplastic nuclei based on chromosomal copy number alterations, and histopathologic characterization of glioma cases. **A-G)** Large scale chromosomal copy number alterations were inferred from RNA expression using InferCNV R package (see methods for details). The heat maps show gains (red) and losses (blue) in case PA1 (**A**), PA2 (**B**), PA3 (**C**), PO1 (**D**), PO2 (two samples – core and margin) (**E**), TB4916 (**F**), and TB4718 (**G**). **H)** Representative Hematoxylin and Eosin-stained section of the brain tissue used for single nuclei RNAseq of the first five cases. Some cases showed clear infiltration with glioma cells PA1, PA2, PA3, and PO2_c, PO2_2. Cases PO1 and PO2_m showed no clear evidence of cellular tumor.

**Figure S3.**
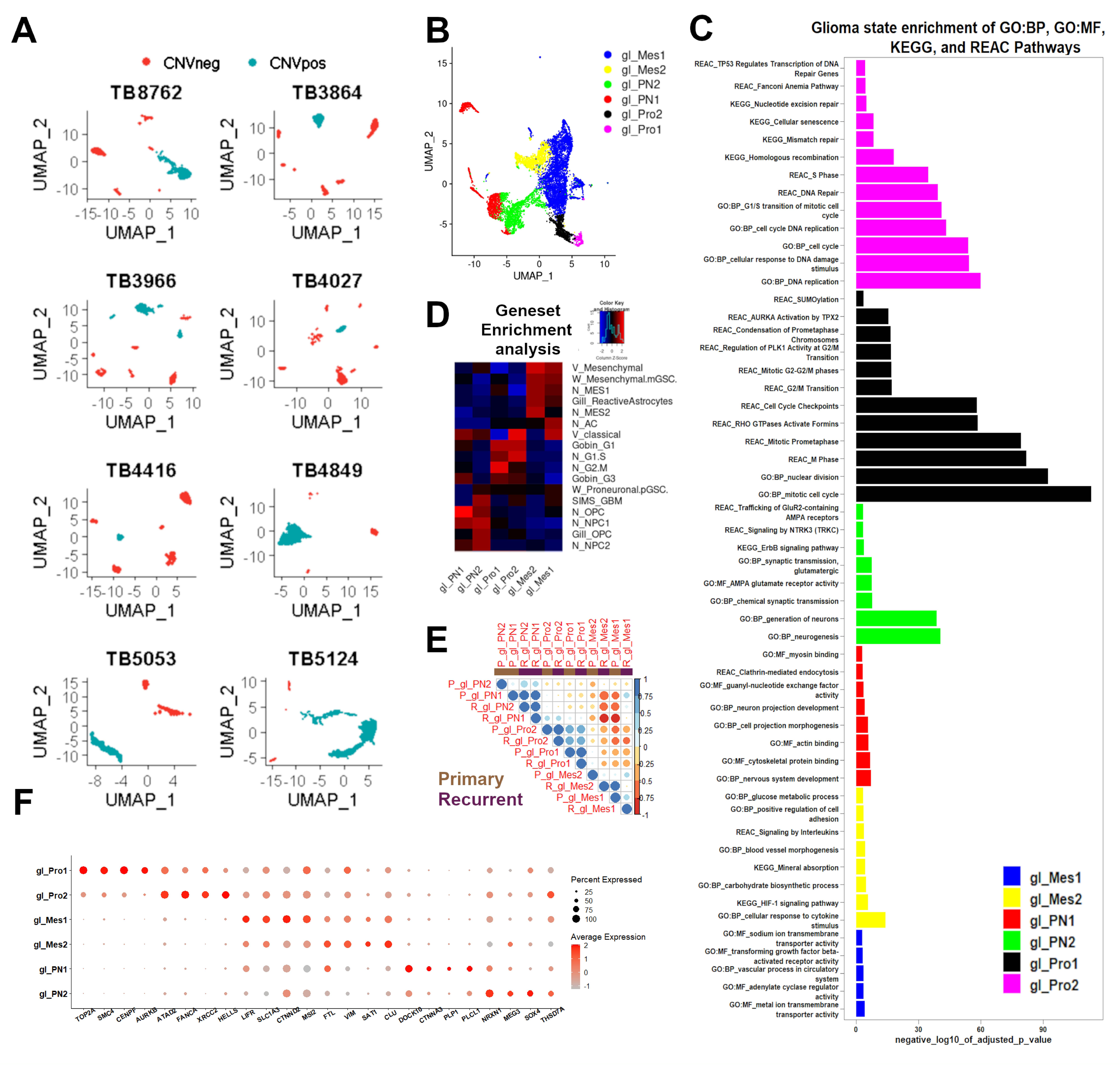
snRNAseq-derived transcriptional states of putative neoplastic nuclei from post-treatment recurrent IDH-wildtype GBM samples. **A**) Uniform-manifold approximation and projection (UMAP) graphs showing putative neoplastic (CNVpos) and non-neoplastic (CNVneg) nuclei from the eight post-treatment recurrent IDH-wildtype glioblastoma cases. **B**) UMAP plot showing all putative CNVpos nuclei from the eight recurrent glioma cases aligned and projected in shared UMAP spaces. The nuclei are color-coded by glioma state: Oligodendrocyte-progenitor-like (proneural - gl_PN1), Neural-progenitor-like (proneural - gl_PN2), Mesenchymal/astrocyte like (gl_Mes1 and gl_Mes2), and proliferative (gl_Pro1 & gl_Pro2). **C**) Gene ontology (GO) term enrichment analysis (KEGG and REACTOME pathways and biological process GO) of the major terms enriched in glioma state top gene markers. The bars represent the negative log10 of the false discovery rate adjusted p.value and are color-coded as in B. **D**) Geneset enrichment analysis (GSEA) of selected genesets from Verhaak et al. 2009, Gobin M et al 2019, Gill et al 2014, and Neftel et al. 2019 showing enrichment of genes specific for states described in the literature in our described glioma states. **E**) Correlation heatmap between glioma states in primary and post-treatment recurrent GBM based on expression on glioma state marker genes. The size and color of the circles denote the strength of correlation. **F**) Gene expression dot plots showing select gene marker expression in glioma states.

**Figure S4:**
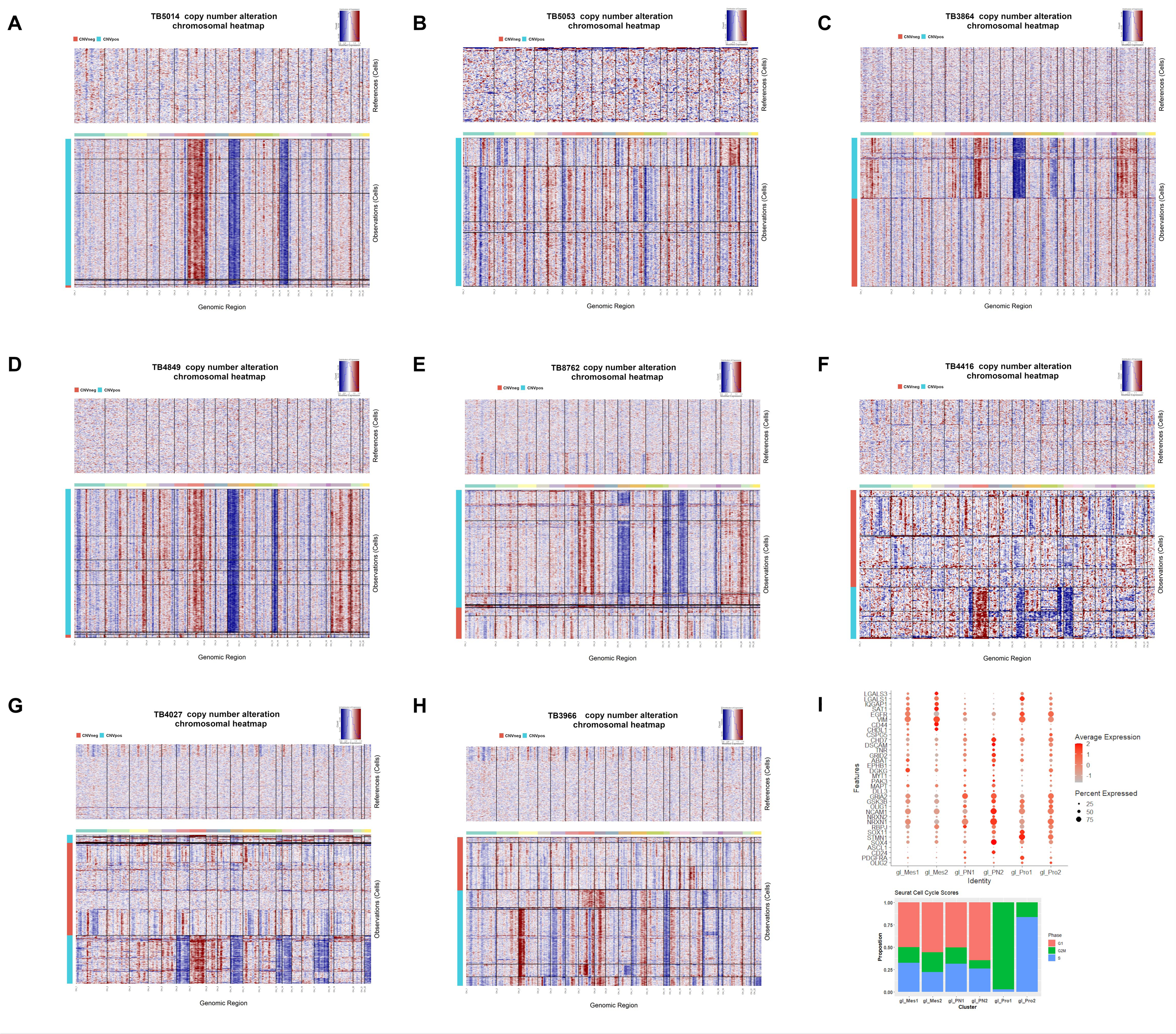
CNV analysis of recurrent glioma samples. **A-H)** Large scale chromosomal copy number alterations were inferred from RNA expression using InferCNV R package. The heat maps show gains (red) and losses (blue) in case TB5014 (**A**), TB5053 (**B**), TB3864 (**C**), TB4898 (**D**), TB8762 (**E**), TB4416 (**F**), and TB4027 (**G**), and TB3966 (**H**). **I**) Dotplot showing expression of select set of markers of both primary and recurrent glioma states. The proportion of each glioma state in cell cycle phases as determined by Seurat cell-cycle scoring is shown on the bottom.

**Figure S5:**
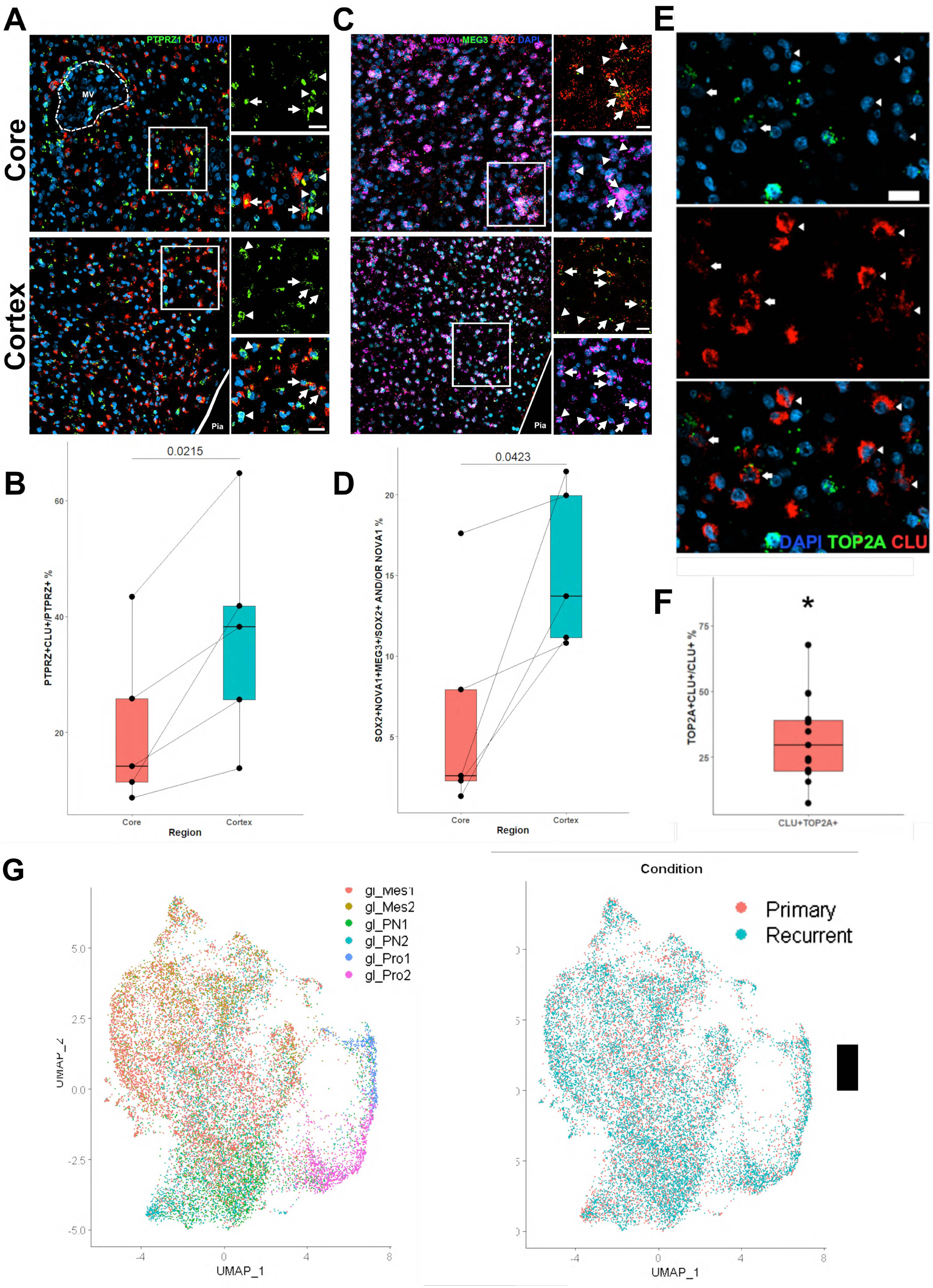
The spatial landscape of glioma states across the cellular tumor and cortex. **A**) Confocal images showing optical sections of in situ hybridization for PTPRZ1 and CLU in the core (upper row) and cortex (lower row). The pial surface is outlined (lower row). High-power images of the insets show that PTPRZ1+ CLU+ cells (arrows) are more abundant in the cortex, while PTPRZ1+CLU- (arrowheads) are more numerous in the core. scale bars = 20 μm. M.V: Microvascular proliferation **B)** Quantification of PTPRZ1 and CLU expression across the core (orange boxplot) and cortex (green boxplot). The data is shown as boxplots, with the bar indicating the median. Paired t-test, N=5 independent samples external to the snRNAseq datasets. The p value is indicated. **C**) Confocal images showing optical sections of in situ hybridization for NOVA, SOX2, and MEG3 in the core (upper row) and cortex (lower row). The pial surface is outlined (lower row). High-power images of the insets show that NOVA1+SOX2+MEG3+ cells (arrows) are more abundant in the cortex, while MEG3-cells (arrowheads) are more numerous in the core. scale bars = 20 μm. **D**) Quantification of MEG3+NOVA1+SOX2+ cells as a proportion of all tumor cells (SOX2+ and/or NOVA1+) across the core (orange boxplot) and cortex (green boxplot). The data is shown as boxplots, with the bar indicating the median. Paired t-test, N=5. The p value is indicated. **E)** Confocal images showing optical sections of in situ hybridization for TOP2A and CLU in the GBM infiltrated tissue. Arrows indicate CLU+TOP2A+ cells, and arrowheads indicate CLU+TOP2A-cells. scale bar = 20 μm. **F)** Quantification of TOP2A and CLU expression. The percentage of TOP2A+CLU+/CLU+ cells is shown as a boxplot. One-sample t-test, N=5 independent samples. *=p value < 0.001. **G)** Integration of primary and recurrent GBM CNVpos nuclei color-coded by glioma state and condition.

**Figure S6:**
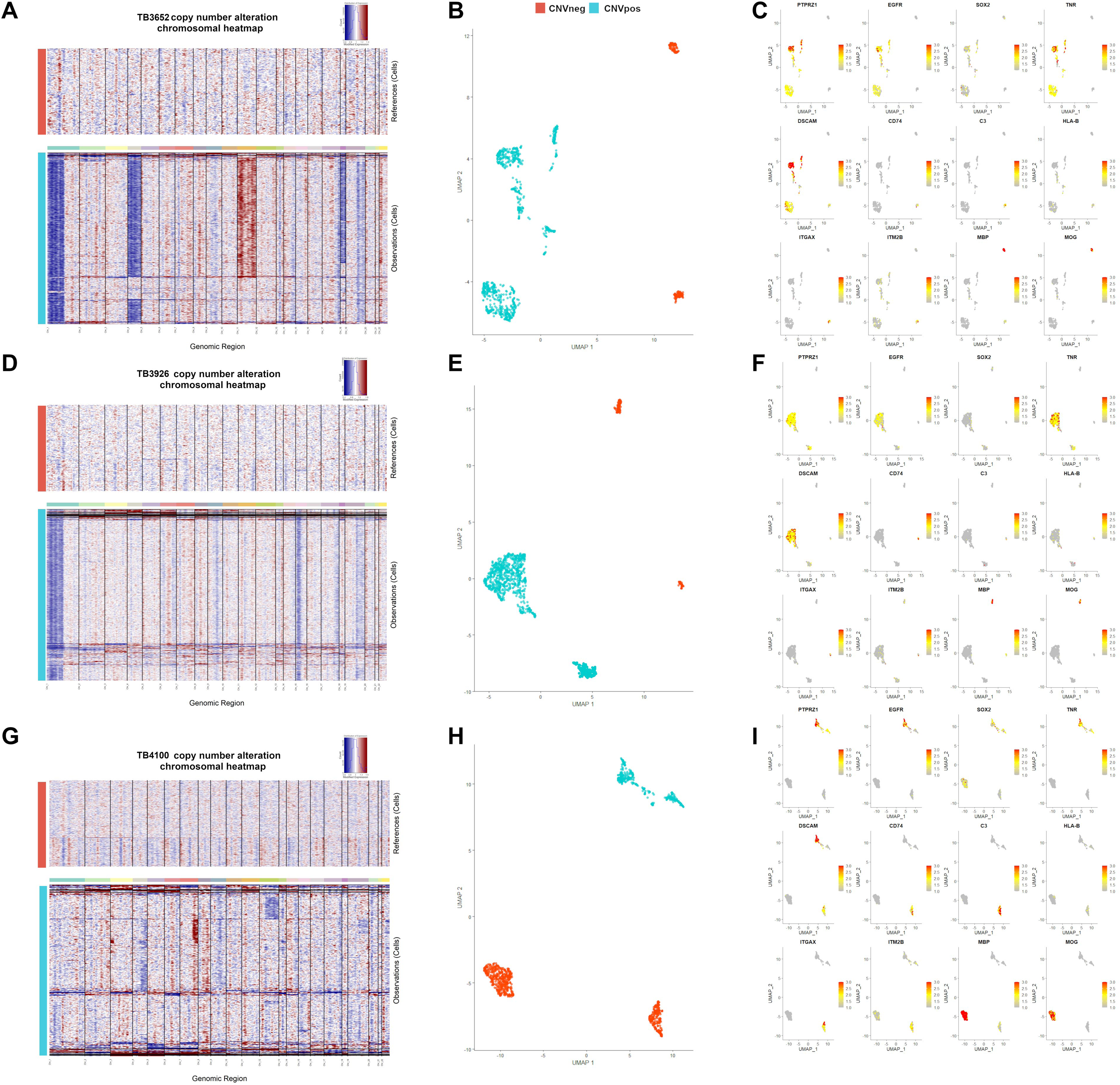
Analysis of Low-grade glioma samples using single nucleus RNAseq. Large scale chromosomal copy number alterations were inferred from RNA expression of cases TB3652 (**A**), TB3926 (**D**) – both IDH1-mutant oligodendrogliomas, and TB4100 (**G**) – IDH-mutant astrocytoma. Uniform manifold approximation and projection (UMAP) plots of the three cases are shown in panels **B**, **E**, and **H**, color-coded by copy number alteration status. Gene expression UMAPs showing markers of tumor cells (PTPRZ1, EGFR, SOX2, TNR, and DSCAM), immune cells (CD74, C3, HLA-B, ITGAX, ITM2B), and oligodendrocytes (MBP, MOG) of cases TB3652, TB3926, and TB4100 in panels **C, F,** and **I**, respectively.

**Figure S7:**
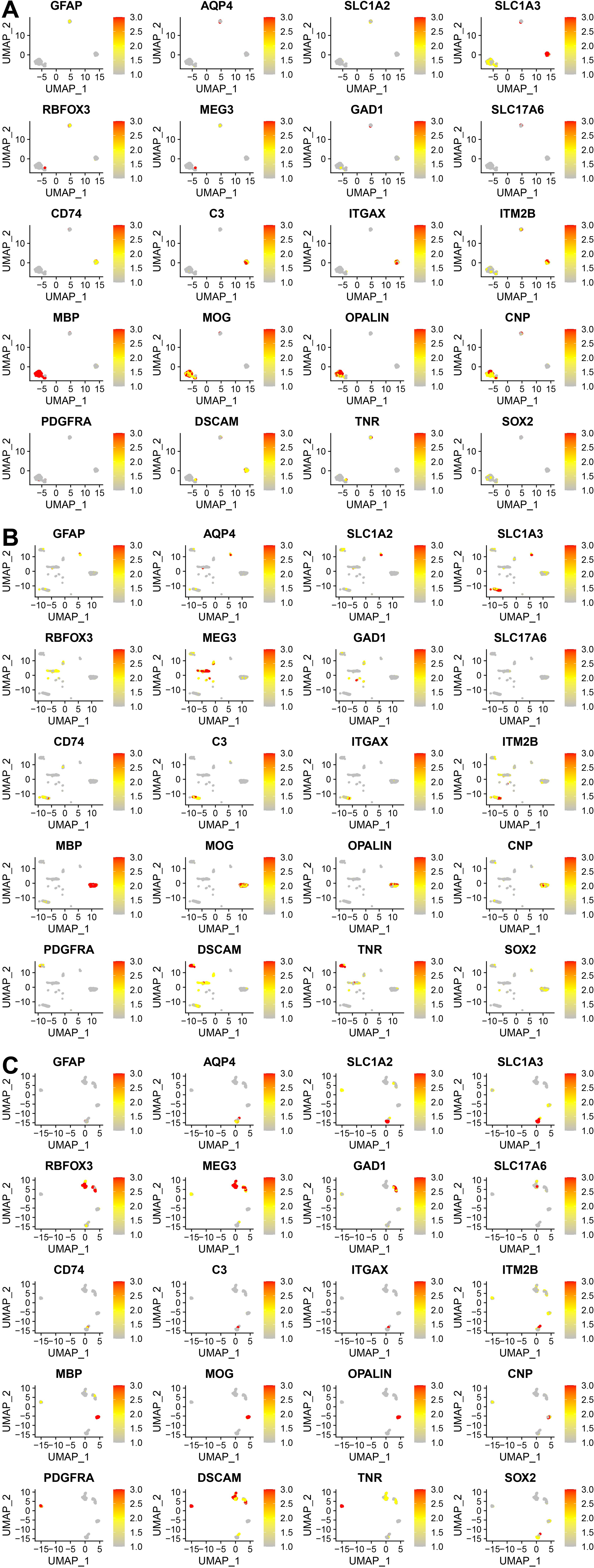
Analysis of Epilepsy samples using single nucleus RNAseq. **A-C**) Uniform-manifold approximation and projection (UMAP) graphs plots showing normalized gene expression of select lineage markers for cases TB4437 (**A**), TB4189 (**B**), TB4957 (**C**). The markers include astrocyte markers (GFAP, AQP4, SLC1A2, and SLC1A3), neuron makers (RBFOX3, MEG3, GAD1, SLC17A6), myeloid markers (CD74, C3, ITGAX, ITM2B), oligodendrocyte markers (MBP, MOG, OPALIN, CNP), and OPC markers (PDGFRA, DSCAM, TNR, and SOX2).

**Figure S8:**
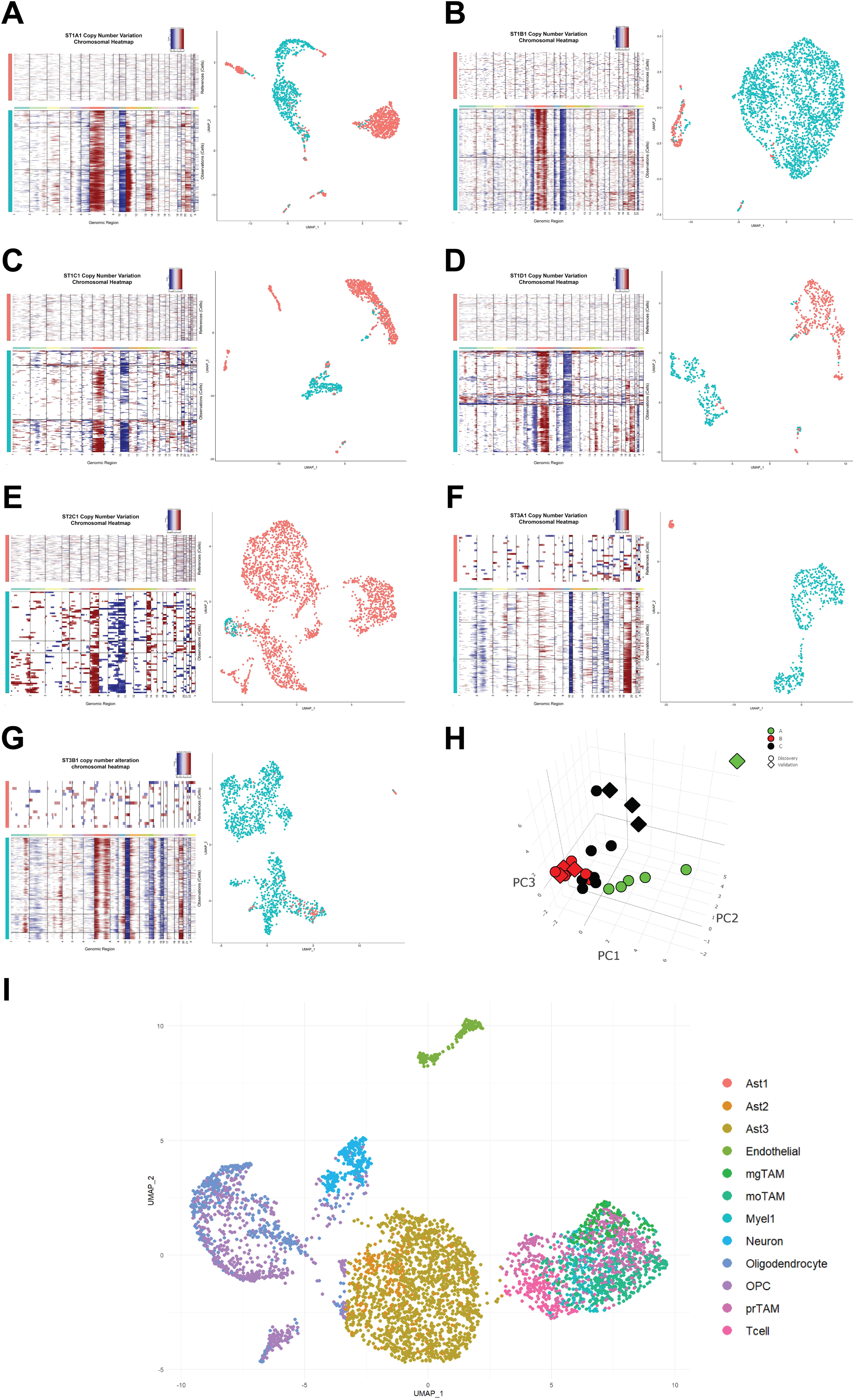
Analysis of Validation single nuclei data set. **A-G)** Heatmaps and UMAP projections of single nuclei extracted from separate sections of tissue that underwent spatial transcriptomics analysis showing copy number variation analysis using the InferCNV R package. Nuclei colored red were classified CNVneg and nuclei colored blue were classified CNVpos. CNVpos and CNVneg cells across samples were integrated and clustered separately before being categorized into specific cell types (see methods.) **H)** The composition of each validation dataset sample was determined and each validation sample was projected onto the PCA axes used to classify the discovery dataset samples. **I)** UMAP projection of all CNVneg nuclei across the validation data set, colored by cell type identity.

**Figure S9:**
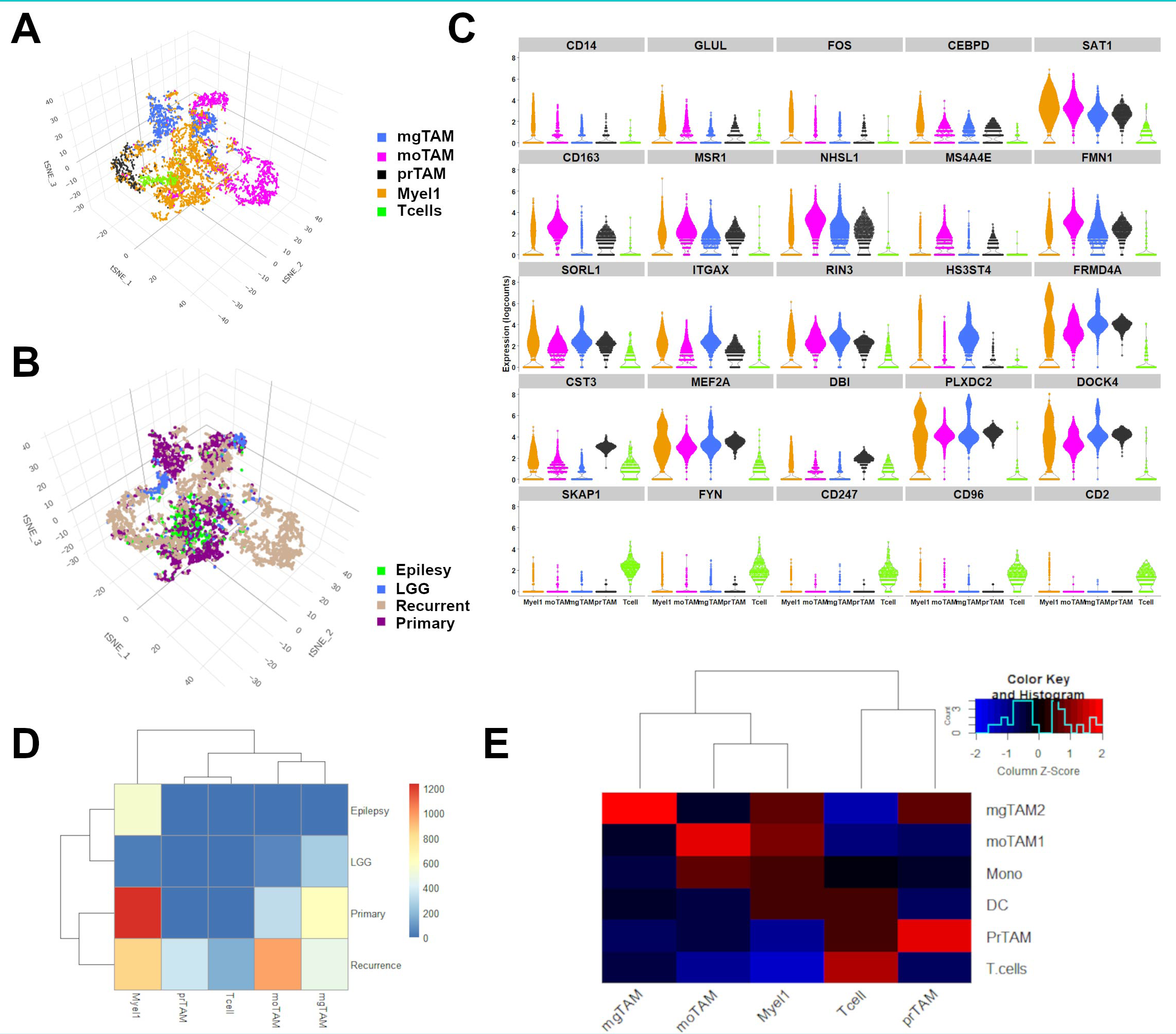
The transcriptional landscape of microglia in glioma. **A**) Uniform-manifold approximation and projection (UMAP) graphs plots showing all myeloid nuclei from color-coded by cluster (**B**) and condition (primary glioma, recurrent glioma, low grade glioma (LGG), and epilepsy (**C**). Gene expression violin plots showing select gene marker expression for the immune cell clusters from top to bottom; Myel1, mgTAM, moTAM, prTAM, and T cells. **D**) Heatmap showing the proportion of nuclei in each cluster (columns) contributed by condition (rows). **E**) Heatmap showing the scaled enrichment scores of gene sets derived from Movahedi et al 2021 in the nuclei pooled from each myeloid cluster.

**Figure S10:**
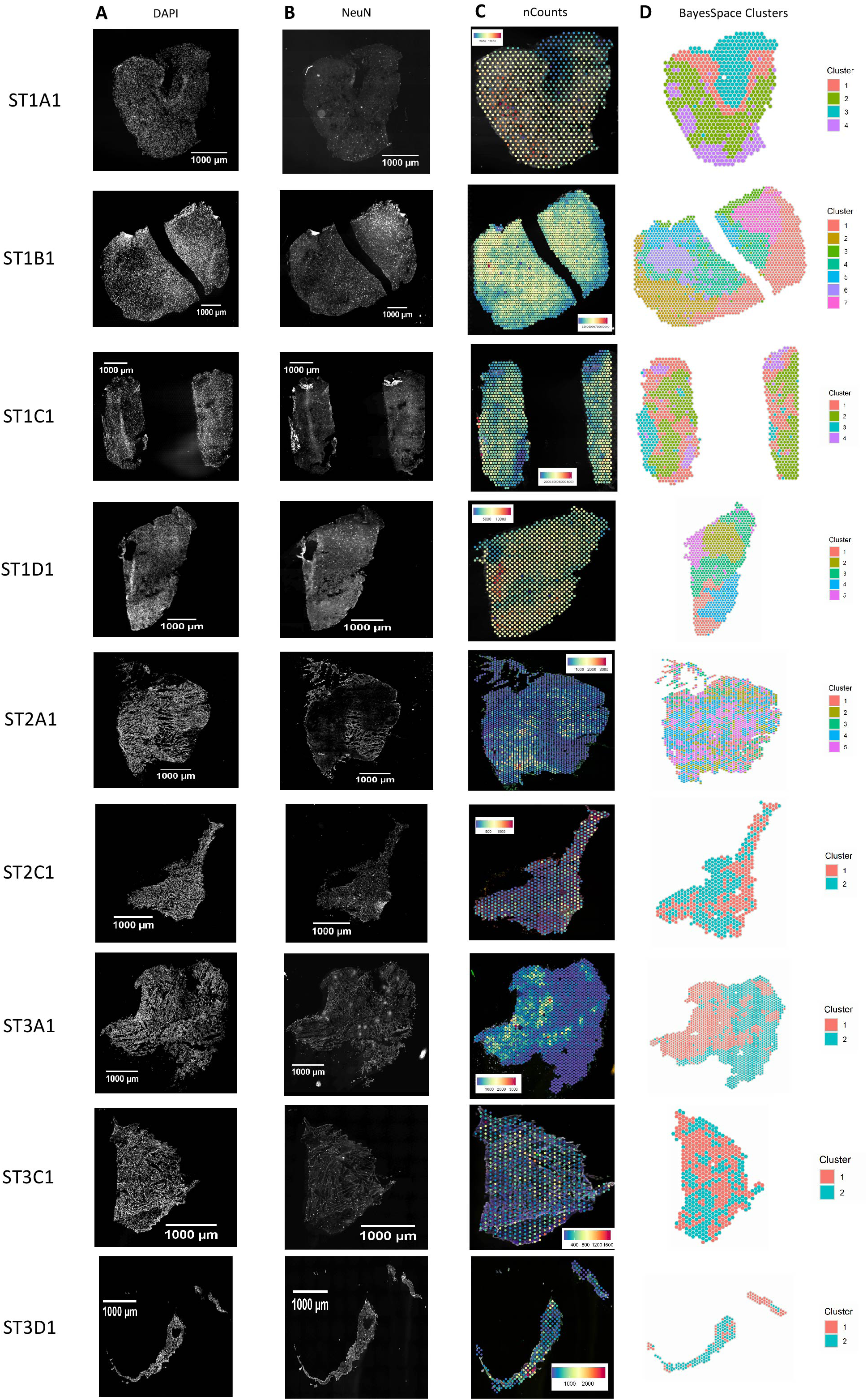
Spatial transcriptomics samples. **A-B)** DAPI and NeuN staining of ST samples. **C)** Spatial transcriptomic images annotated with number of unique genes observed at each spot. **D)** BayesSpace-generated clusters overlaid on each ST sample. The number of clusters for each sample was determined through maximization of the modified Bayesian Information Criterion (MBIC).

**Figure S11:**
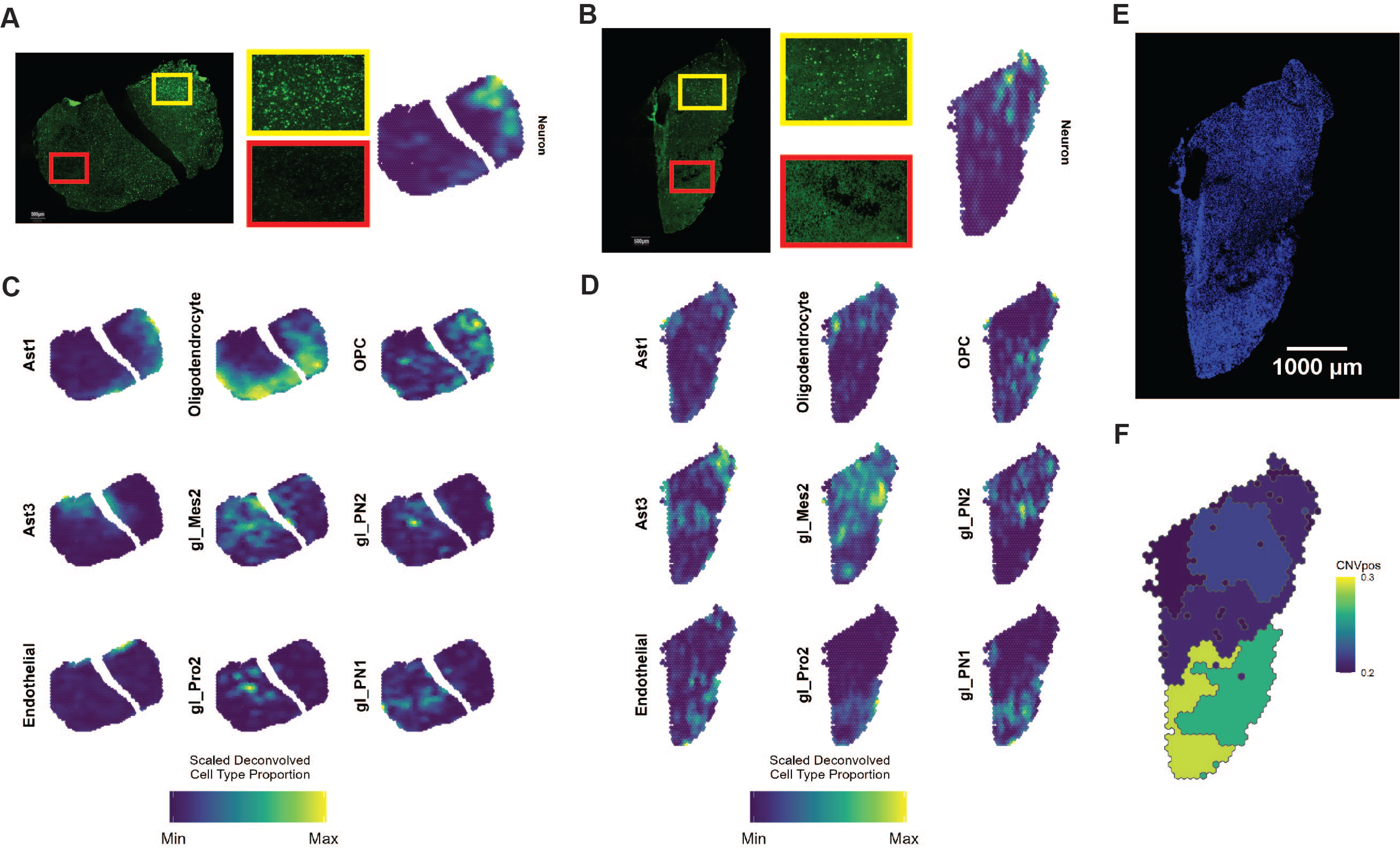
Deconvolution of Spatial Transcriptomic Samples. **A-B)** Representative images of staining for NeuN alongside the deconvolved proportions of Neurons in samples ST1B1 and ST1D1. Red and yellow insets show detailed view of NeuN staining that correlates with the patterns of heterogeneity shown by the deconvolved proportion of neurons. **C-D)** Deconvolved proportions of selected cell types that comprise each tissue state projected onto maps of samples ST1B1 and ST1D1 respectively. **E)** Sample ST1D1 stained for DAPI and **F**) sample ST1D1 segmented by BayesSpace into clusters and shaded based on the proportion of CNVpos cell types determined by deconvolution. Clusters with a higher density of nuclei were correlated with a higher proportion of CNVpos cell types across the dataset.

**Figure S12:**
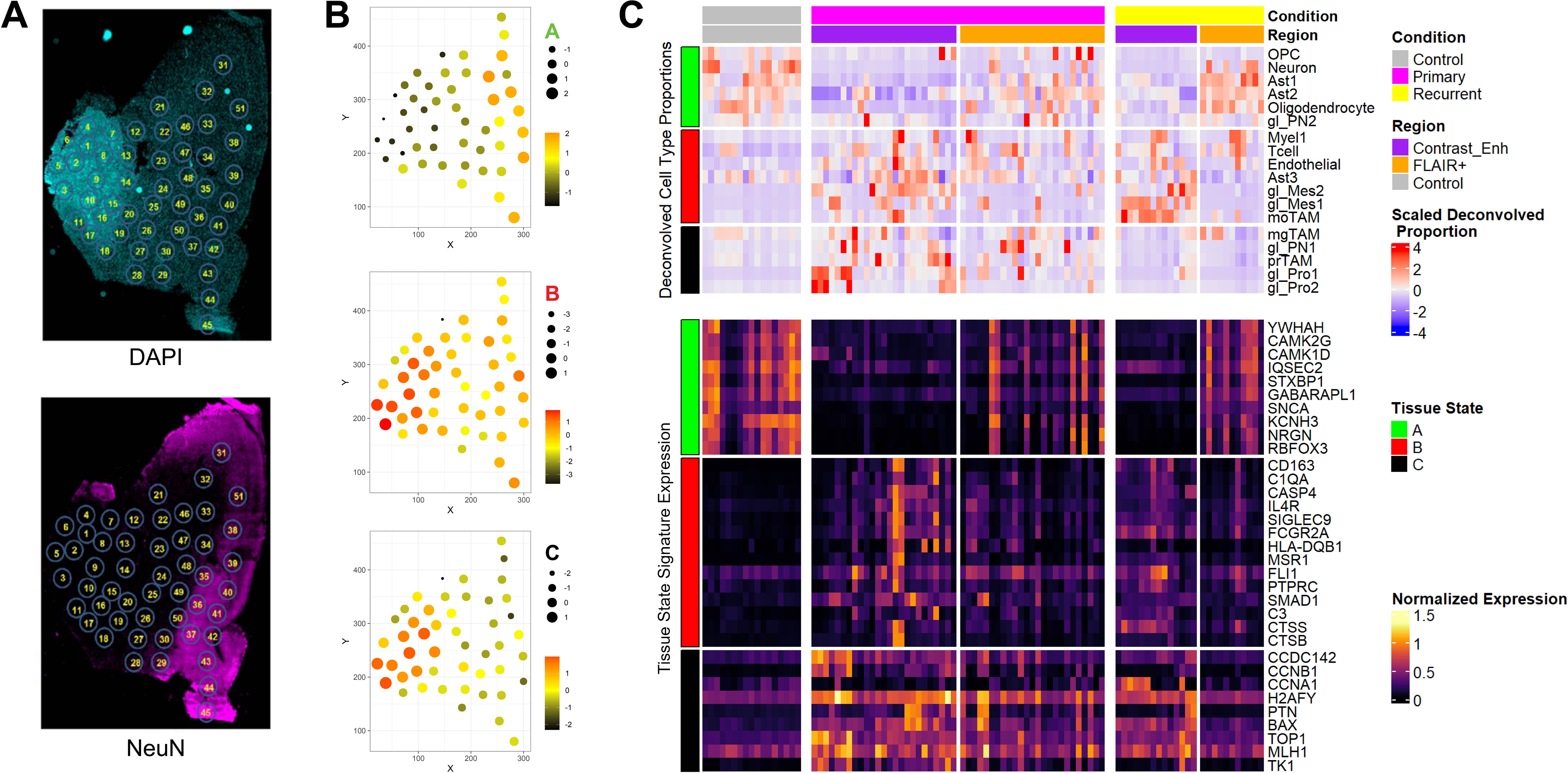
The spatial landscape of glioma margins. **A**) Outline of spatial transcriptomic analysis of infiltrating GBM. DAPI (left) and NeuN (right) immunostains of frozen sections from case PO2, for which snRNAseq was done. Each circle represents a biopsy on which bulk RNAseq was done. After the biopsies were taken, the specimen was bisected along the dashed white line (y-axis) and subjected to snRNASeq. **B**) Enrichment analysis of each of the spatially mapped biopsies using the genesets of the three compositional clusters (see text for details) displaying normalized single sample GSEA enrichment scores for the tumor cluster (C - upper panel), the tumor-reactive cluster (B– middle panel), and the normal brain cluster (A – lower panel). The enrichment scores are coded by color and size. The normalized RNA data for the spatial biopsy map is available in an interactive web interface at https://vmenon.shinyapps.io/gbm_expression/. **C**) Heatmaps showing the scaled proportion of 18 cell types obtained by deconvolution and corresponding normalized expression of markers from tissue-state signatures for each sample as applied to the Gill et al. 2014 MRI localized biopsy dataset (n=92). Results are stratified by condition and by MRI localization. Cell types/markers are annotated with their corresponding tissue-states on the left.

**Figure S13:**
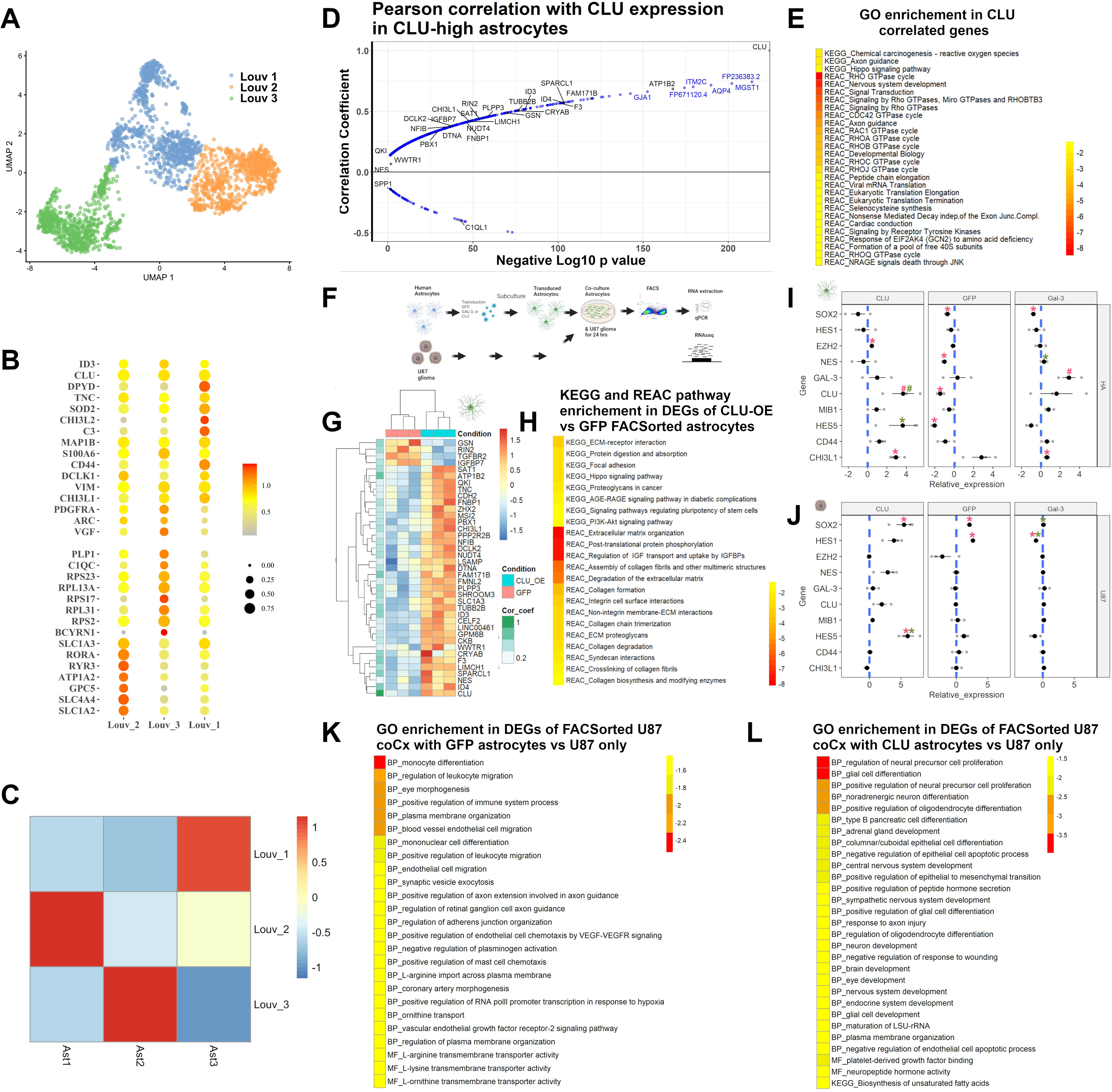
Astrocytes influence glioma gene expression. **A)** UMAP projection of astrocytes clustered using Louvain clustering on shared-nearest neighbor graphs created using igraph – k=500. Three clusters are shown. **B)** Select cluster markers shown in dotplots. **C**) Scaled overlap between identities of astrocytes designated using clustering on geneset enrichment (described in figure 1) versus unbiased clustering described in **A** of this figure - scaled by column. **D**) Correlation analysis showing the Pearson correlation coefficients (y-axis) of genes that correlate with CLU expression in CLU-high astrocytes – defined as having normalized expression in the third or fourth quantiles. The p value is shown on the x-axis. **E**) Gene ontology enrichment analysis of the genes that positively correlate with CLU from A. **F**) Outline of astrocyte glioma co-culture experiment. The negative log10 of the p value of enrichment is indicated. **G**) Heatmap of normalized gene expression of astrocytes with CLU over-expression (CLU_OE) versus GFP control astrocytes co-cultured with U87 glioma cells. The genes are annotated by the correlation coefficient with CLU from A. **H**) KEGG and Reactome pathway enrichment in the genes differentially expressed in FACSorted CLU_OE astrocytes versus GFP astrocytes after co-culture with U87 glioma. The negative log10 of the p value of enrichment is indicated. **I-J**) Real-time quantitative PCR of select genes from CLU_OE, GFP, or LGALS3_OE astrocytes (**I**) and U87 glioma (**J**) FACSorted from co-culture experiment. The genes were selected based on relevance to Ast3 signature or genes relevant to glioma biology). The Log normalized delta-delta Ct values are shown on the y-axis. The p values are indicated. #: one-tailed paired t-test, *: two-tailed paired t-test. N = 3 independent FACsorting experiments. Error bars indicated standard error of the mean. The comparison group: control astrocytes not co-cultured with glioma for “red” * or #; control astrocytes co-cultured with U87 cells for “green” * or #. **K-L**) Gene ontology enrichment analysis of genes differentially expressed in U87 cells co-cultured with GFP astrocytes (versus U87 cells not co-cultured **K**) or co-cultured with CLU_OE astrocytes versus U87 glioma co-cultured with GFP astrocytes (**L**). The negative log10 of the p value of enrichment is indicated.

